# p38β/MAPK11 Deficiency Exacerbates Cardiac Structural and Electrophysiological Remodeling and Contributes to Immune Dysregulation in the Aging Heart

**DOI:** 10.64898/2026.06.18.733182

**Authors:** Katy A. Trampel, Batool Salman, Lara Leoni, Sophie Green, Noor Saleem, Ava Adli, Daniele Procissi, Igor R. Efimov, Tatiana Efimova

## Abstract

Aging is a major risk factor for cardiac diseases, including heart failure, myocardial infarction, and arrhythmias. Activation of p38 MAPKs regulates cardiac remodeling and contributes to age-related cardiac dysfunction. However, the isoform-specific roles of p38 kinases in the aging heart remain poorly understood. Although p38β has been reported to exert cardioprotective effects in models of doxorubicin-induced cardiotoxicity and ischemia-reperfusion, its role in cardiac aging remains unclear. Here, we investigated the role of p38β using p38β germline knockout (p38β^-/-^) mice. Aged p38β^-/-^ mice exhibited increased LV hypertrophy, QT prolongation, calcium mishandling, heightened susceptibility to arrhythmias, increased myocardial fibrosis, and an altered inflammatory microenvironment, compared with age-matched wild-type controls. Transcriptomic profiling revealed that p38β deletion reprograms the cardiac transcriptome in aged mice, suppressing innate immune and proteostasis-related pathways while promoting adaptive immune activation, developmental, extracellular vesicle-mediated, and ion-transport pathways. Collectively, these findings identify p38β as a critical regulator of structural, electrophysiological, and immune homeostasis in the aging heart and demonstrate that its loss promotes maladaptive remodeling and arrhythmogenic vulnerability.

**NEW AND NOTEWORTHY:** We identify p38β as a previously unrecognized regulator of cardiac aging. Systemic loss of p38β disrupts structural, electrophysiological, and immune homeostasis in the aging heart, revealing its protective role in maintaining cardiac function with age. These findings underscore the importance of isoform-specific p38 signaling and suggest that broadly targeting p38 MAPKs may have unintended consequences in age-related cardiovascular diseases.

## INTRODUCTION

The global population is living longer, with individuals over 80 projected to reach 137 million worldwide by 2050 (1). Advanced age is one of the strongest independent risk factors for cardiovascular disease (CVD), even in the absence of overt myocardial injury. Aging is associated with heart failure, atrial fibrillation, ventricular tachyarrhythmias, and sudden cardiac death (1–3). Among these, ventricular tachycardia (VT) and fibrillation (VF) are major contributors to morbidity and mortality in age-related CVDs (4). The increased frequency of ventricular arrhythmias with age reflects progressive remodeling of the myocardial substrate. Cardiac aging is a gradual process, not an abrupt switch, marked by changes in calcium handling, ion channel expression and activity, repolarization, intercellular coupling, and extracellular matrix remodeling (4,5). These alterations in the electrophysiological substrate can promote triggered activity, reentry, and conduction heterogeneity, beginning in midlife and increasing the risk of arrhythmias as age advances (1,5,6).

Despite the well-established association between chronological aging and ventricular arrhythmias, the pathophysiological mechanisms linking age-related myocardial stress to arrhythmogenesis remain incompletely defined, limiting the development of targeted therapies. In addition to electrophysiological remodeling, the aging heart exhibits increased oxidative stress, mitochondrial dysfunction, DNA damage, and chronic inflammation, all of which elevate myocardial stress and electrical instability. This age-induced pathological remodeling activates intracellular stress-activated signaling pathways, including p38 mitogen-activated protein kinases (MAPKs), which are crucial regulators of adaptive and maladaptive stress responses in the aging heart (7–9).

The mammalian p38 MAPK family comprises four isoforms: p38α, p38β, p38γ, and p38δ, encoded by the *MAPK14*, *MAPK11*, *MAPK12*, and *MAPK13* genes, respectively. Although these isoforms share high sequence homology, they exhibit distinct functional and regulatory mechanisms (10). Historically, p38α/p38β and p38γ/p38δ were grouped based on sequence similarity (10). However, growing evidence supports isoform-specific signaling mechanisms and non-redundant functions under myocardial stress. In particular, a recent study reported activation of the p38γ and p38δ isoforms in the hearts of aged mice compared with young controls, and simultaneous overexpression of activated forms of these kinases in the heart promoted stress-induced ventricular arrhythmogenesis (7). In contrast, the roles of the p38β isoform in cardiac electrophysiology and age-related cardiac electrical remodeling remain poorly understood.

In the heart, p38β has been reported to play a protective role in ischemia/reperfusion injury by regulating oxidative stress and apoptosis in vitro and in vivo (9,11–14). In addition, our recent findings showed that germline deletion of p38β worsened doxorubicin-induced cardiotoxicity (DIC) in female mice (14). Notably, DIC is known to mimic key features of accelerated cardiac aging (8). Therefore, we explored whether p38β might play a protective role under conditions of age-related cardiac stress. Here, we demonstrate that germline p38β knockout exacerbates age-related cardiac structural and electrical remodeling, contributes to immune dysregulation and inflammaging, and induces global transcriptomic changes in the aging heart, thereby promoting maladaptive remodeling and increasing vulnerability to ventricular arrhythmogenesis.

## METHODS

All animal protocols were approved by the Institutional Animal Care and Use Committee at Northwestern University and comply with the National Institutes of Health Guide for the Care and Use of Laboratory Animals.

### Mice

Mice used in this study were on a C57BL/6 background. Wild-type (WT) mice were bred in-house or purchased from Jackson Laboratory (stock no. 000664). The generation of mice with a germline deletion of p38β was previously reported (15). p38β knockout mice (p38β^-/-^) are viable and fertile, with no overt abnormalities. Genotyping was performed as described (15). We used age-matched groups of male and female mice aged 13.5 to 20 months. The WT and p38β^-/-^experimental groups were also sex-matched, as sex distribution did not differ significantly between the groups (p = 0.89, Fisher’s exact test). All mice were housed in a pathogen-free environment under a 12-hour dark/light cycle at 70-74°F with 30–70% humidity and were provided food and water ad libitum.

### Electrocardiography (ECG)

Conscious ECGs were recorded using the emka Technologies ecgTUNNEL device (emka Technologies, Sterling, VA). WT mice and p38β^-/-^ mice were guided into the ecgTUNNEL, where their paws were restrained and positioned in contact with the silver electrodes. After a 5-minute acclimation period, ECG signals were recorded using leads I and II. ECG parameters were measured using a custom MATLAB code to quantify P-wave, PR interval, QRS duration, QT interval, corrected QT (QTc), and RR interval.

### Echocardiography

WT mice and p38β^-/-^ mice were anesthetized with 2% isoflurane in oxygen and maintained at a heart rate above 350 beats per minute during imaging. The animals were placed in a supine position on a heated imaging platform at 37°C, with anesthesia depth monitored. Chest hair was removed using depilatory cream, and ultrasound gel was applied. LV short-axis images were acquired with a 40-MHz probe from the high-frequency ultrasound system (Prospect, S-Sharp, New Taipei City, Taiwan). LV short-axis images were analyzed using Prospect Software.

### Langendorff Perfusion

Aged WT mice (7 males and 8 females) and p38β^-/-^ mice (4 males and 5 females) were deeply anesthetized with 5% isoflurane in oxygen. Anesthesia depth was confirmed by the absence of the toe pinch reflex, after which cervical dislocation was performed. Hearts were rapidly excised following thoracotomy, and the aorta was cannulated and connected to a Langendorff perfusion system. Hearts were retrogradely perfused with warmed (37°C) and oxygenated (95% oxygen and 5% carbon dioxide) modified Tyrode’s solution (in mM, NaCl 130, NaHCO_3_ 24, NaH_2_PO_4_ 1.2, MgCl_2_ 1, glucose 5.6, KCl 4, and CaCl_2_ 1.8) at pH 7.4. Perfusion flow rate was maintained at 1.0 – 2.0 mL/min and adjusted as needed to achieve a coronary pressure of approximately 80 mmHg. Pseudo-ECGs were recorded via electrodes placed in the superfusion bath.

### Optical Mapping

Langendorff-perfused hearts were electromechanically uncoupled with 15 μM blebbistatin (Cayman Chemicals 13186) and equilibrated for 20 minutes. Hearts were then loaded and perfused with the voltage-sensitive dye RH237 (30 µL of 1 mg/mL RH237 stock solution plus 970 µL Tyrode’s solution, Biotium 61018) and the calcium fluorescent dye Rhod2-AM (30 µL of 1 mg/mL Rhod2-AM stock solution plus 30 µL of Pluronic F-127 plus 940 µL Tyrode’s solution, Thermo Fisher Scientific R 1244 and Biotium 59055, respectively). Both dyes were delivered via coronary perfusion. Hearts were illuminated using a 520 ± 5 nm excitation light source (Prizmatix UHP-Mic-LED-520), and fluorescence emission was collected using CMOS cameras (MiCAM, SciMedia). Baseline recordings were obtained to determine the pacing capture threshold, after which hearts were paced at 1.5 times the capture threshold amplitude using a platinum epicardial point electrode. Data were analyzed using Rhythm 3.0 from optical action potentials (V_m_ RT, APD_30_, APD_50_, APD_80_, CV_L_, CV_T_, and AR_CV_) and optical calcium transients (Ca^2+^ RT, CaTD_30_, CaTD_50_, CaTD_80_, and Ca^2+^ τ).

### Arrhythmia Inducibility

Cardiac arrhythmia inducibility was assessed using burst pacing protocols and administration of 300 nM isoproterenol. Pseudo-ECGs were recorded to detect and analyze arrhythmias in ex vivo hearts. Arrhythmias were classified into two types: VF and VT, and by duration. VF was identified by irregular ECG waves with high variability and no discernible morphological ECG wave parameters. VT was characterized by an increased heart rate and regular, wide-complex waves. Arrhythmias lasting more than 4 seconds but less than 20 seconds were considered non-sustained; those longer than 20 seconds were classified as sustained. The incidences of VT and VF, along with total arrhythmia events and their durations, were recorded from pseudo-ECG data and optical mapping. Early after-depolarizations (EADs) were identified through optical action potential (OAP) traces as secondary depolarizations occurring during phase 2 or 3 of repolarization before complete membrane repolarization.

### Mouse Cytokine/Chemokine Array

WT and p38β^-/-^ male mice at 17 months of age (n = 4 per group) were anesthetized with an isoflurane/oxygen mixture. Anesthesia depth was confirmed by loss of the pedal reflex. Mice were euthanized by cervical dislocation followed by thoracotomy. The aorta was cannulated, and the heart was flushed with PBS. Ventricular tissue was isolated and flash-frozen in liquid nitrogen. The tissue was homogenized in RIPA lysis buffer (Sigma 89901) containing protease and phosphatase inhibitors (Thermo Fisher A32955). Total protein concentration was measured using the Pierce BCA kit (Thermo Fisher 23227) according to the manufacturer’s instructions. Cytokine and chemokine levels were quantified using the Mouse Cytokine/Chemokine 68-Plex Discovery Assay® Array (MD68) from Eve Technologies.

### Picrosirius Red Staining

Hearts from WT and p38β^-/-^ male and female mice at 17 and 20 months of age, respectively (n = 4 per group), were embedded in Tissue-Tek® O.C.T. Compound (VWR 25608-930) and cryopreserved by gradual freezing in isopentane cooled with liquid nitrogen. Samples were stored at -80°C until processing. Hearts were sectioned transversely. For Picrosirius Red staining, three LV images were acquired at 20x magnification (Zeiss) and averaged per sample. Fibrosis was quantified using a custom MATLAB-based color deconvolution algorithm, and Picrosirius Red-positive pixels per total area were reported as the percent fibrosis.

### Wheat Germ Agglutinin (WGA) Staining

Transverse sections of hearts from WT and p38β^- /-^ male and female mice at 17 and 20 months of age, respectively (n = 4 per group), were fixed in 4% PFA for 30 minutes, washed in PBS, and incubated with WGA-Alexa Fluor (Thermo Fisher W11261) for 2 hours. Sections were washed and mounted with ProLong™ Glass Antifade Mountant (Thermo Fisher P36984). Six non-overlapping left ventricular fields per sample were imaged at 10× magnification (Nikon) in regions with circular cardiomyocyte profiles. Twenty cardiomyocyte cross-sectional areas per image were measured using FIJI, yielding a total of 120 cells per heart. Values were averaged to obtain a single mean cross-sectional area per sample.

### Bulk RNA Sequencing

Hearts from WT and p38β^-/-^ male mice at 17 months of age (n = 4 per group) were perfused with PBS and preserved in RNALater^TM^ (Millipore Sigma MFCD03453003). Bulk RNA sequencing of ventricular tissue was performed by Genewiz (Azenta Life Sciences). Sequencing reads were aligned to the mouse genome using STAR, and transcript abundance was quantified with RSEM. Differential gene expression analysis was conducted using DESeq2 with apeglm shrinkage; significance was set at padj < 0.1. Principal component analysis was performed on regularized log-transformed data. Differentially expressed genes (DEGs) were defined as padj < 0.1 and |log2FC| > 1 (FC, fold change). DEGs were visualized with volcano plots. Gene Ontology (GO) Biological Processes (BP), Cellular Components (CC), and Molecular Function (MF), and Kyoto Encyclopedia of Genes and Genomes (KEGG) pathway analyses were conducted using enrichGO and enrichKEGG, respectively, at padj < 0.1. For Gene Set Enrichment Analysis (GSEA), genes were ranked by log2FC and analyzed against Hallmark gene sets (MSigDB, Category H) using clusterProfiler, with q < 0.05 considered significant. Data are available in the NIH Gene Expression Omnibus (GEO) repository under accession number GSE328163.

### Micro-Computed Tomography (micro-CT)

Micro-CT imaging was performed at the Center for Translational Imaging Small Animal Molecular Imaging Core Facility (RRID: SCR_017878) on 17-month-old WT (n = 4), p38β^+/-^ (n = 3), and p38β^-/-^ (n = 4) male mice to assess bone morphology. Images were acquired on a Mediso NanoScan microCT (Mediso, Hungary) and analyzed using VivoQuant (Perceptive, USA) and ITK-SNAP software (16). X-ray settings were 70 kilovoltage peak (kVp), 280 μA, 300 ms exposure, and 720 projections. For femoral analysis, left distal femoral metaphyseal trabecular bone volume fraction (BV/TV) was quantified at 20 µm resolution by selecting a metaphyseal region distal to the growth plate and calculating trabecular bone volume relative to total tissue volume. Cortical bone thickness was measured in both left and right femurs at the midshaft by averaging four measurements taken at standardized anterior, posterior, and lateral positions on cross-sectional images. For cranial analysis, skull scans were acquired at 30 µm resolution. Interparietal trabecular bone BV/TV was quantified by manually defining the region of interest.

### Statistical Analysis

All data were analyzed using GraphPad Prism 9 (GraphPad Software, Inc.). Data are presented as mean ± standard deviation. Sample sizes were determined a priori using a power analysis with α = 0.05 and β = 0.8. No animals or samples were excluded from analysis. To ensure that the WT and p38β^-/-^ experimental groups were sex-matched, Fisher’s exact test was performed on the counts of male and female mice within each genotype group. Comparisons between two independent groups were performed using unpaired two-tailed Student’s t-tests, including analyses of cytokine and chemokine profiling, ECG parameters, and echocardiographic measurements. For comparisons among more than two groups, one-way ANOVA was used with genotype as the independent variable. Post hoc comparisons were performed using unpaired Student’s *t*-tests with Tukey correction to evaluate differences in bone volume. For unpaired binary parameters, such as arrhythmia and EAD incidence, Fisher’s exact test was used. For parameters exhibiting restitution properties, including V_m_ RT, APD_30_, APD_50_, APD_80_, CV_L_, CV_T_, AR_CV_, Ca^2+^ RT, CaTD_30_, CaTD_50_, CaTD_80_, and Ca^2+^ τ, data were fit using an exponential plateau model. Best-fit parameters were compared between groups using the sum-of-squares F test. Statistical significance was defined as p < 0.05.

## RESULTS

### p38β deletion promotes LV structural and mechanical remodeling in aged mice

We assessed LV structure and function using LV short-axis M-mode echocardiography in age- and sex-matched groups of aged WT and p38β^-/-^ male and female mice (Fig. 1A-1R and Supplemental Table 1). Mice ranged from 13.5 to 19.75 months of age, with a mean of approximately 16 months in the WT and p38β^-/-^ experimental groups (Fig. 1A and 1C). Body weight did not differ significantly between the groups, with mean body weights of 32.4 g for WT and 33.4 g for p38β^-/-^ groups (Fig. 1D). However, LV mass normalized to body weight was higher in aged p38β^-/-^ mice compared to WT controls (Fig. 1E), indicating hypertrophic remodeling. Representative LV short-axis M-mode echocardiograms show differences in anterior wall structure between groups (Fig. 1F). LV functional parameters, including ejection fraction (EF), fractional shortening (FS), stroke volume (SV), and cardiac output (CO), along with structural parameters, such as LV anterior wall thickness during systole and diastole (LVaWs and LVaWd), LV posterior wall thickness during systole and diastole (LVpWs and LVpWd), and LV end diameter during systole and diastole (LVESD and LVEDD), as well as LV end volume during systole and diastole (LVESV and LVEDV), were measured (Fig. 1G – 1R). Aged p38β^-/-^ mice showed an increase in SV compared to their WT counterparts (Fig. 1I), indicating increased blood volume ejected per heartbeat. Consistent with this mechanical remodeling, aged p38β^-/-^ mice showed increased LVaWs and LVaWd (Fig. 1K and 1L). Additionally, LVEDV was trending upward in the p38β^-/-^ group compared with WT controls (Fig. 1R). Echocardiographic analyses were also stratified by sex, and no sex-dependent differences were observed (Supplemental Fig. 1, Supplemental Tables 2 and 3).

**Fig. 1.**
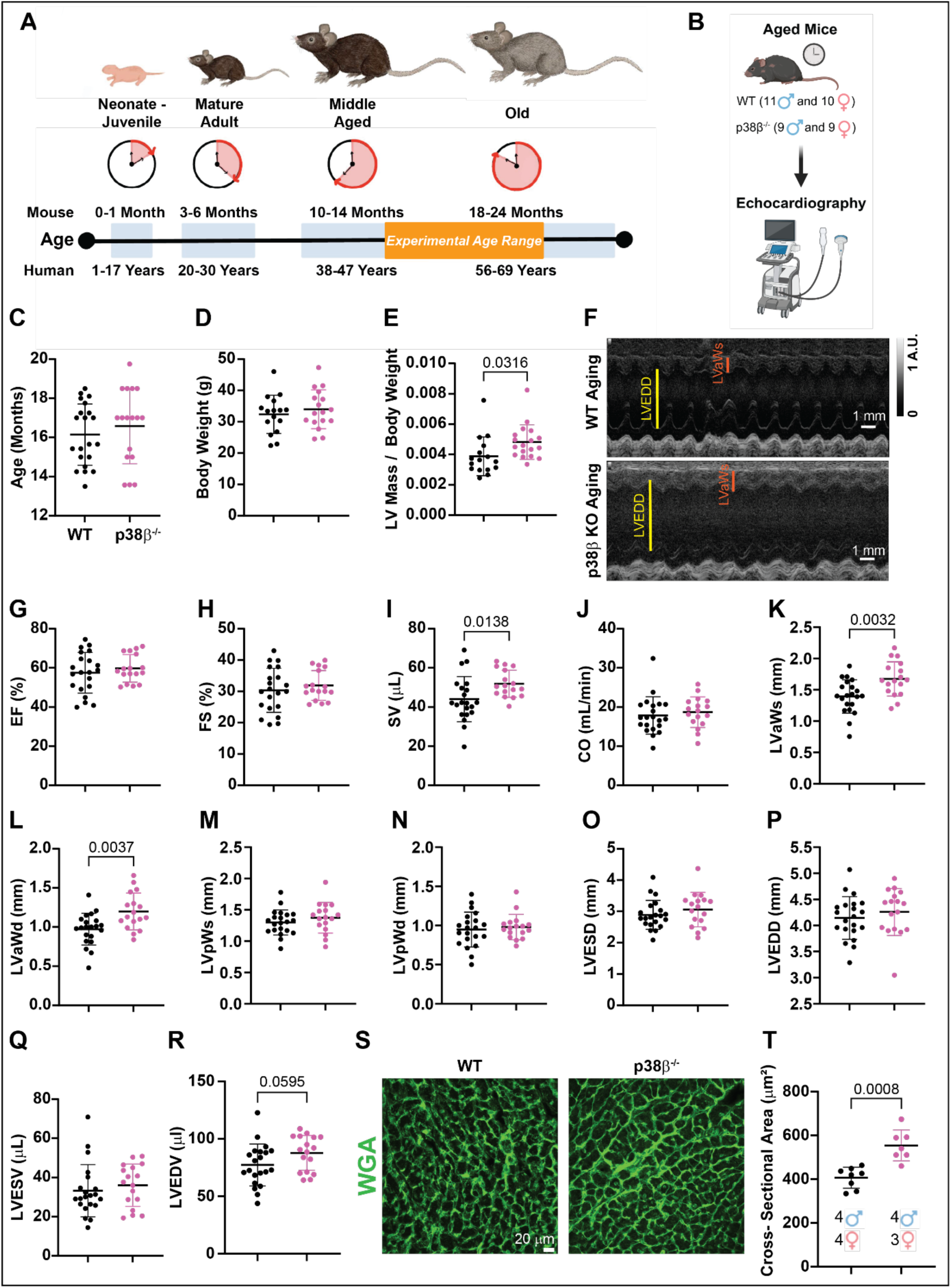
p38β deficiency increases LV mass, SV, LVaWs, LVaWd, LVEDV, and cardiomyocyte cross-sectional area in aged mice. *A:* Schematic illustrating life phases in mice and their correspondence to those in humans (adapted from the white paper “Aged C57BL/6J Mice for Research Studies” by The Jackson Laboratory). The age range of mice used in this study is shaded in orange. *B:* Schematic illustrating echocardiography acquisition in aged WT and p38β^-/-^ mice. The experimental groups included male and female mice and were sex-matched, as sex distribution did not differ significantly between the WT and p38β^-/-^ cohorts (p = 0.89, Fisher’s exact test). *C:* Age distribution of mice used in this study. *D:* Body weights. *E:* LV mass measured via echocardiography relative to body weight. *F:* Representative echocardiograms from aged WT and p38β^-/-^ hearts. *G – P:* Quantification of structural and mechanical parameters from echocardiograms in the specified groups. *S:* Representative images of wheat germ agglutinin (WGA) staining (green) in LV sections from 17-20-month-old WT and p38β^-/-^ mice. *T:* Cardiomyocyte cross-sectional area quantified in WGA-stained hearts. Comparisons between groups were assessed via an unpaired Student’s *t*-test to determine statistical significance. Ejection fraction (EF), fractional shortening (FS), stroke volume (SV), cardiac output (CO), LV anterior and posterior wall thicknesses during systole and diastole (LVaWs, LVaWd, LVpWs, LVpWd), LV end-systolic diameter (LVESD), LV end-diastolic diameter (LVEDD), and LV volume during systole and diastole (LVESV and LVEDV). Sample sizes for age, body weight, and echocardiography: n = 21 in the WT group; n = 18 in the p38β^-/-^ group. Sample sizes for WGA staining: n = 8 in the WT group; n = 7 in the p38β^-/-^ group.

To assess whether these structural changes were associated with cardiomyocyte hypertrophy, wheat germ agglutinin (WGA) staining was used to quantify cardiomyocyte cross-sectional area. Representative images of WGA-stained LV sections from aged WT and p38β^-/-^ male and female mouse hearts are shown in Fig. 1S. Quantitative analysis revealed that cardiomyocyte cross-sectional area was increased in aged p38β^-/-^ mice compared with their WT counterparts (Fig. 1T), indicating cardiomyocyte hypertrophy. Collectively, these data demonstrate that p38β deficiency promotes both structural and mechanical LV remodeling in the aging heart.

### p38β deficiency promotes QT and QTc prolongation in aged mice

Global cardiac electrical function was assessed by recording conscious ECGs in aged WT and p38β^-/-^mice, as illustrated in Fig. 2A and summarized in Supplemental Table 1. ECG wave parameters were quantified, including P wave, PR interval, QRS duration, QT interval, RR interval, and QTc (Fig. 2B). Representative morphological differences in ECG waveforms between genotypes showed prolonged ventricular repolarization in aged p38β^-/-^ mice compared with WT controls (Fig. 2C). Consistent with these findings, aged p38β^-/-^ mice exhibited prolonged QT and QTc intervals (Figs. 2G and 2I) compared with their WT counterparts, reflecting delayed ventricular depolarization and repolarization. No differences in P wave duration, PR interval, QRS duration, or RR interval were observed (Figs. 2D-2H). Furthermore, sex-stratified analyses recapitulated the genotype-dependent differences observed in the combined analysis, with both aged p38β^-/-^males and females exhibiting QT and QTc interval prolongation relative to their corresponding WT controls (Supplemental Fig. 2, Supplemental Tables 2 and 3). To determine whether these electrophysiological alterations persist with advanced aging, ECGs were recorded in a separate cohort of 20-month-old female WT and p38β^-/-^ mice (Supplemental Fig. 3). Consistent with findings in middle-aged mice, 20-month-old female p38β^-/-^ mice also demonstrated QT and QTc prolongation (Supplemental Figs. 3D and 3F). These findings demonstrate that p38β deficiency causes maladaptive electrical remodeling characterized by QT prolongation in the aging heart, supporting a cardioprotective role for p38β during aging.

**Fig. 2.**
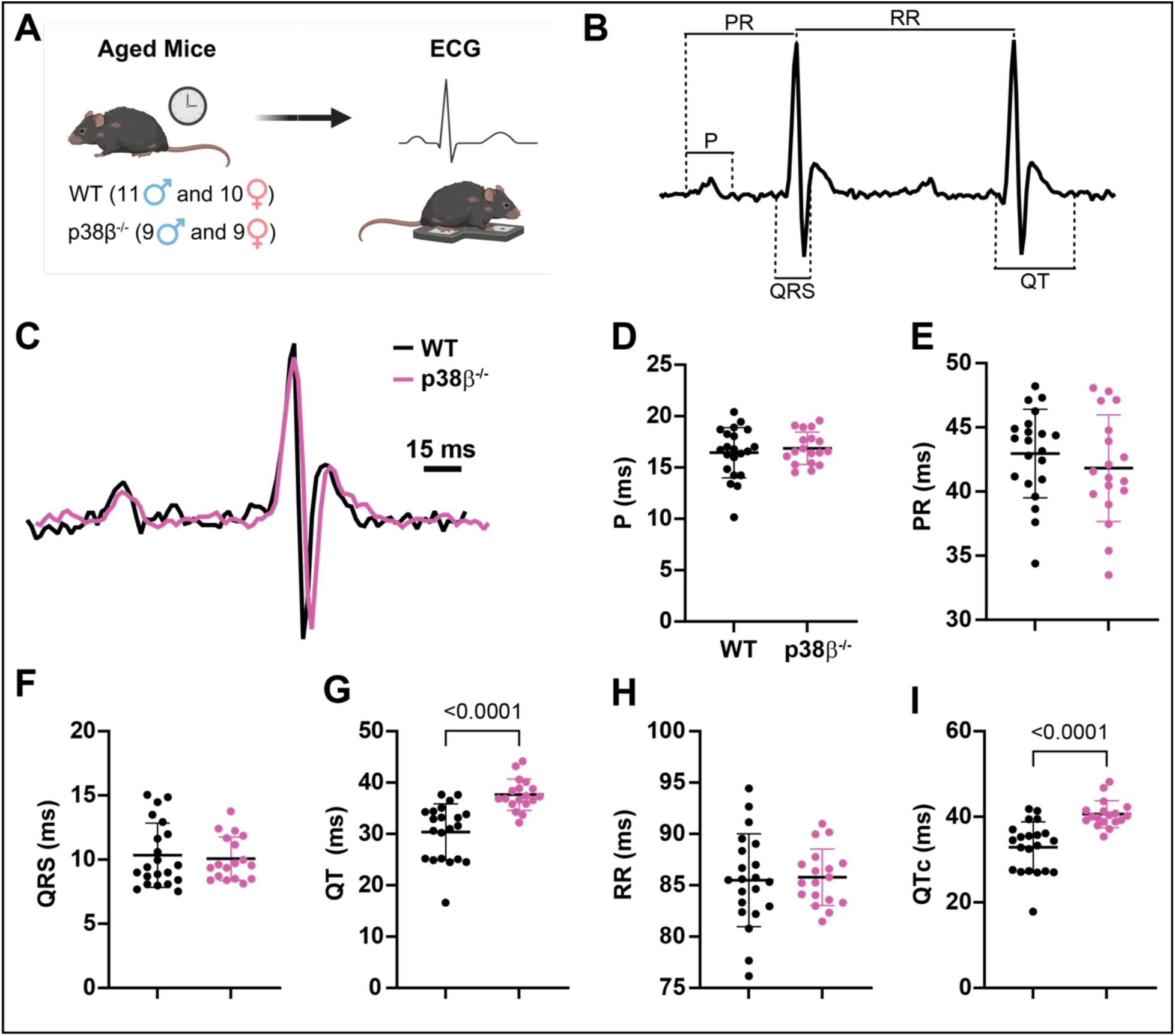
p38β deficiency prolongs QT and QT_C_ in aged mice. *A:* Schematic of ECG acquisition in aged WT and p38β^-/-^ mice. *B:* Representative ECG traces with annotated waveform parameters. *C:* Representative ECGs from WT (black) and p38β^-/-^ (magenta) mice. *D – I:* Quantification of P wave, PR interval, QRS duration, QT interval, RR interval, and QT_C_ interval. Unpaired Student’s *t*-tests were used to assess statistical significance between groups. Sample sizes: n = 21 in the WT group; n = 18 in the p38β^-/-^ group.

**Fig. 3.**
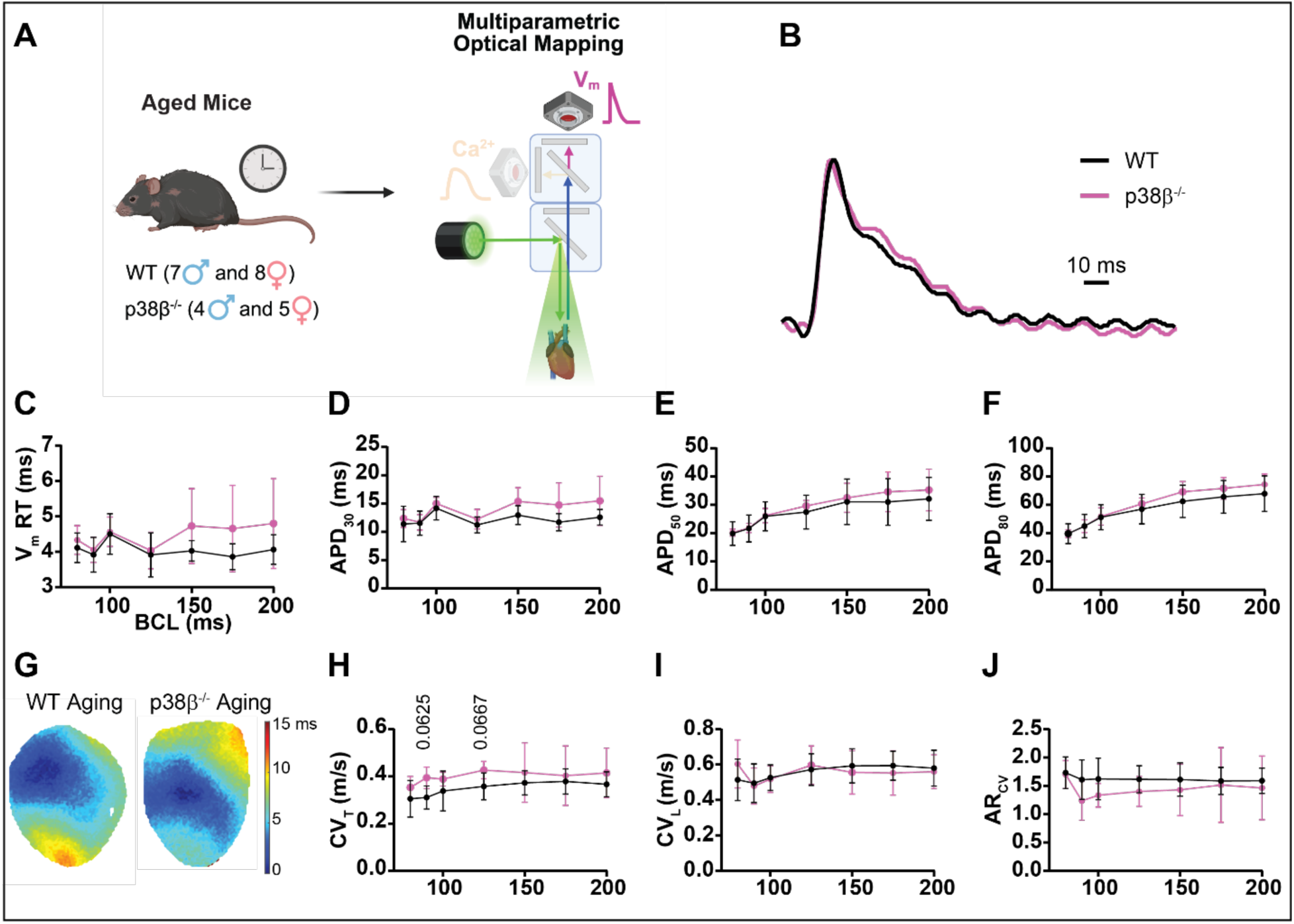
p38β deficiency does not significantly alter transmembrane potential in aged murine hearts ex vivo. *A:* Schematic of ex vivo multiparametric optical mapping of aged WT and p38β^-/-^ murine hearts. *B:* Representative optical action potentials recorded from aged WT and p38β^-/-^ hearts. *C – F:* Quantification of transmembrane potential rise time (V_m_ RT) and action potential duration at 30% (APD_30_), 50% (APD_50_), and 80% (APD_80_) repolarization. *G:* Representative activation maps for the specified groups. *H – J:* Quantification of CV_T_, longitudinal (CV_L_) conduction velocity, and anisotropic ratio of conduction velocity (AR_CV_). Nonlinear regression with an exponential plateau model was used to assess differences among groups. BCL, basic cycle length. Sample sizes: n = 15 in the WT group; n = 9 in the p38β^-/-^ group.

### p38β deficiency does not significantly affect ventricular depolarization or repolarization in aged murine hearts ex vivo

Ex vivo optical mapping of transmembrane potential (V_m_) and intracellular calcium transients (CaT) was performed in aged WT and p38β^-/-^hearts from both male and female mice (Fig. 3A). Representative OAP traces from aged WT and p38β^-/-^ hearts ex vivo showed similar morphology, with no differences in depolarization or repolarization (Fig. 3B). To evaluate electrophysiological remodeling, we measured OAP restitution parameters, including V_m_ rise time (V_m_ RT), action potential duration at 30% (APD_30_), 50% (APD_50_), and 80% (APD_80_), longitudinal conduction velocity (CV_L_), transverse conduction velocity (CV_T_), and anisotropic conduction velocity (AR_CV_). Across a range of basic cycle lengths (BCLs), no differences were observed in OAP restitution properties between aged WT and p38β^-/-^hearts (Figs. 3C – 3J). A trend toward increased CV_T_ was observed in p38β^-/-^ hearts at shorter cycle lengths (Fig. 3H). Collectively, these findings suggest that p38β deficiency does not significantly affect ventricular depolarization or repolarization in aged hearts ex vivo.

### p38β deficiency promotes calcium mishandling in aged murine hearts ex vivo

Using ex vivo multiparametric optical mapping, we assessed intracellular Ca^2+^ flux in aging WT and p38β^-/-^ murine hearts (Fig. 4A). Representative optical Ca^2+^ transient (CaT) traces from aged p38β^-/-^ hearts showed altered waveform morphology compared with WT controls (Fig. 4B). To evaluate excitation-contraction coupling (ECC) remodeling, we measured CaT restitution parameters, including calcium rise time (Ca^2+^ RT), CaT duration at 30% (CaTD_30_), 50% (CaTD_50_), and 80% (CaTD_80_) reuptake, and calcium decay (Ca^2+^ τ). As shown in Figs. 4C – 4E, aged p38β^-/-^hearts had prolonged Ca^2+^ RT, CaTD_30_, and CaTD_50_ compared with WT controls. In contrast, no differences were observed in CaTD_80_ or Ca^2+^ τ between genotypes (Figs. 4F, 4G). Overall, these findings suggest that p38β deficiency impairs calcium release and early calcium reuptake in aged hearts.

**Fig. 4.**
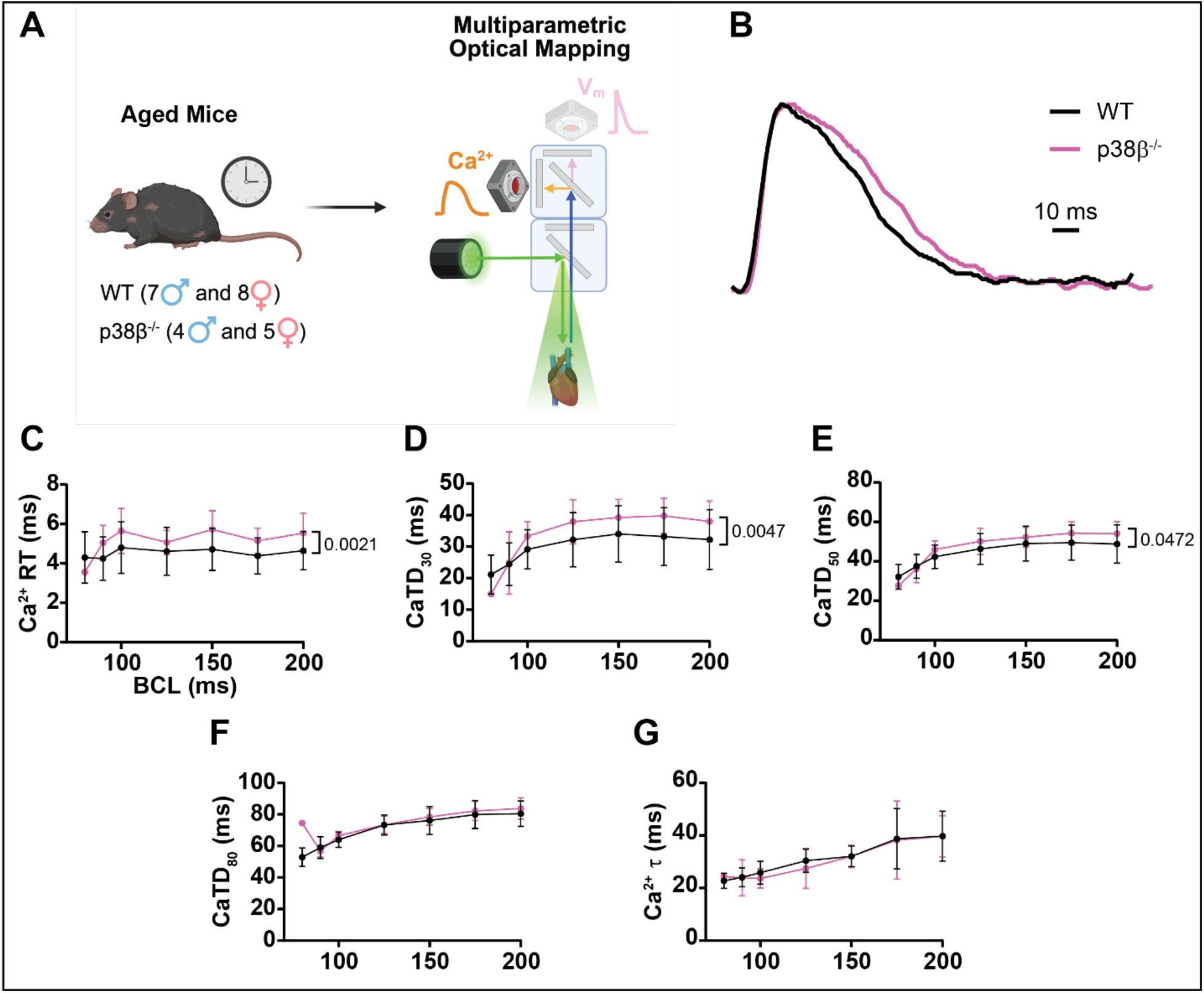
p38β deficiency prolongs Ca^2+^ transient rise time (Ca^2+^ RT), calcium transient duration at 30% (CaTD_30_), and 50% (CaTD_50_) reuptake in aged murine hearts ex vivo. *A:* Schematic of ex vivo multiparametric optical mapping of aged WT and p38β^-/-^ murine hearts. *B:* Representative optical Ca^2+^ transient recorded from aged WT and p38β^-/-^ hearts. *C – G:* Quantification of Ca^2+^ RT, CaTD_30_, CaTD_50_, calcium transient duration at 80% (CaTD_80_) reuptake, and Ca^2+^ decay (Ca^2+^ 1). Nonlinear regression with an exponential plateau model was used to assess differences among groups. BCL, basic cycle length. Sample sizes: n = 15 in the WT group; n = 9 in the p38β^-/-^ group.

### p38β deficiency promotes arrhythmogenesis in aged murine hearts ex vivo

Next, ventricular arrhythmia inducibility was assessed ex vivo using rapid pacing and β-adrenergic stimulation (Fig. 5A). Representative pseudo-ECG recordings illustrate sinus rhythm, ventricular tachycardia (VT), ventricular fibrillation (VF), and non-sustained arrhythmias (Figs. 5B – 5E). Burst pacing and isoproterenol administration induced VT and VF in both aged WT and p38β^-/-^ hearts. Ventricular arrhythmia inducibility and duration were quantified from pseudo-ECG recordings. As shown in Fig. 5F, p38β deficiency increased ventricular arrhythmia inducibility, with all aged p38β^-/-^ hearts analyzed exhibiting VT/VF episodes. In contrast, no differences in arrhythmia duration were observed between genotypes (Fig. 5G).

**Fig. 5.**
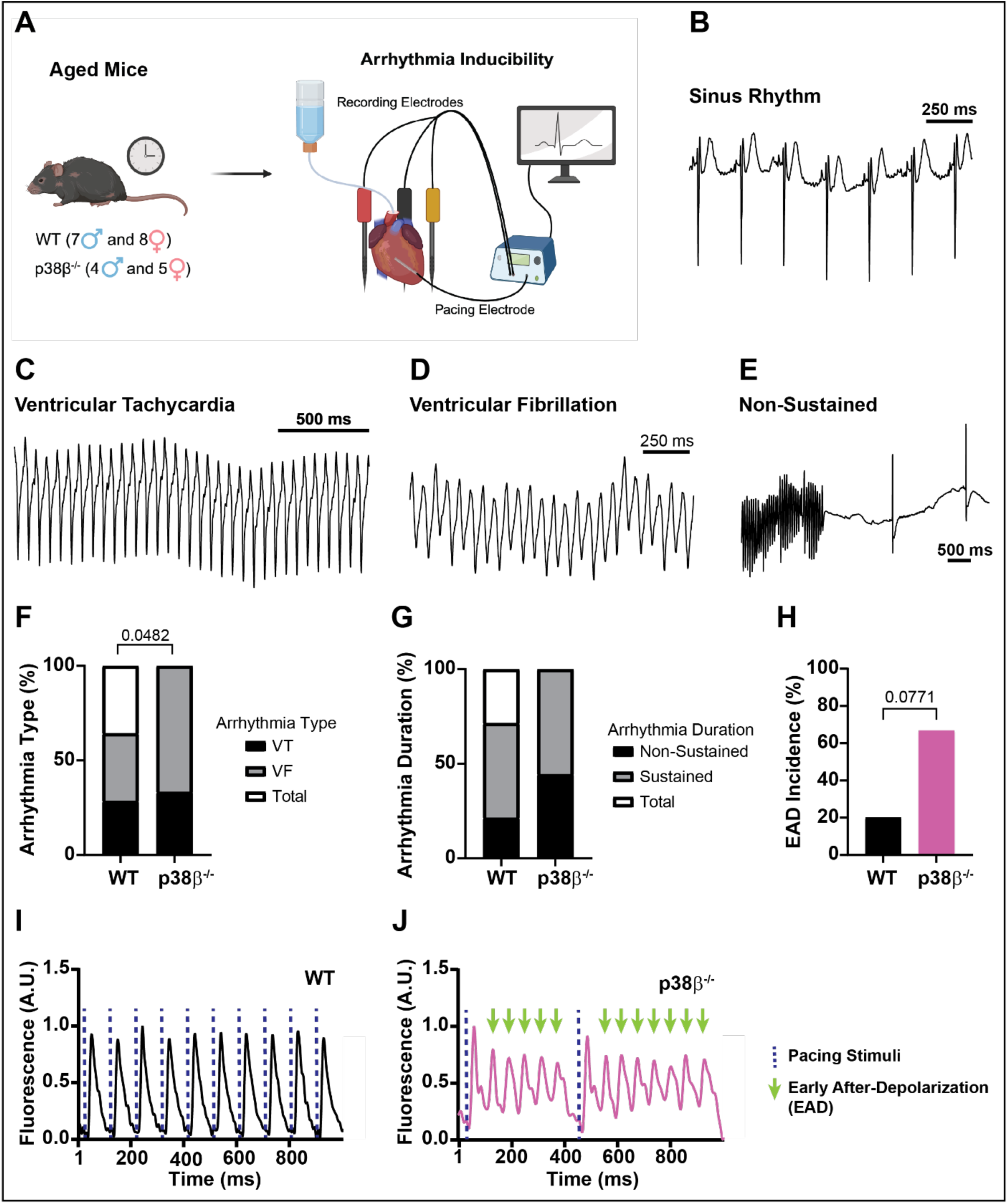
p38β deficiency promotes ventricular arrhythmias and a trend toward increased EAD incidence in aged ex vivo murine hearts. *A – D:* Representative recordings of Sinus Rhythm, Ventricular Tachycardia (VT), Ventricular Fibrillation (VF), and Non-Sustained arrhythmia pseudo-ECG. *E:* Quantification of VT, VF, and total arrhythmia incidence. *F:* Quantification of non-sustained (5 < time < 20 s), sustained (time > 20s), and total duration incidence. *G, H:* Optical action potentials showing paced beats (blue dashed line) and EADs (green arrows). *I:* Quantification of EAD incidence. A Fisher’s exact test was performed to compare groups. Sample sizes: n = 15 in the WT group; n = 9 in the p38β^-/-^ group.

Given the increased arrhythmia inducibility in aged p38β^-/-^ hearts, we next assessed the incidence of early afterdepolarizations (EADs), a known trigger of ventricular arrhythmias. Representative OAP traces from aged WT hearts show paced OAPs (Fig. 5I), whereas OAPs from aged p38β^-/-^ hearts exhibited spontaneous depolarizations during the repolarization of paced beats, consistent with EADs (Fig. 5J). As shown in Fig. 5H, aged p38β^-/-^ hearts showed a trend toward increased EAD incidence (p = 0.0771) compared with WT controls. Thus, p38β deficiency increases ex vivo ventricular arrhythmia susceptibility in aged murine hearts, potentially through enhanced triggered activity.

### p38β deficiency promotes cardiac inflammation and inflammaging in aged mice

Aging is associated with a chronic, low-grade inflammatory state called inflammaging, which contributes to age-related cardiac dysfunction (17). Systemic and cardiac inflammation contribute to the development of various arrhythmias, including atrial fibrillation, long-QT syndrome, Torsades de Pointes, and atrioventricular conduction block (18).

To determine whether p38β loss affects age-related myocardial inflammation, comprehensive cytokine and chemokine profiling was performed by assessing the expression of 68 proinflammatory biomarkers in hearts isolated from 17-month-old WT and p38β^-/-^ male mice (Figs. 6A – 6D, Supplemental Table 4). Notably, levels of B-cell activating factor (BAFF), a component of the senescence-associated secretory phenotype (SASP) linked to immunosenescence, inflammaging, and adverse cardiovascular outcomes, were increased in hearts from aged p38β^-/-^ mice compared with their WT counterparts (Fig. 6E) (19). Additionally, levels of interleukin-17A (IL-17A), a proinflammatory cytokine involved in adverse pro-fibrotic cardiac remodeling and arrhythmogenesis and another key mediator of inflammaging (20), were higher in aged p38β^-/-^ hearts compared to WT controls (Fig. 6K). Moreover, aged p38β^-/-^ hearts also showed trends toward increased levels of the inflammaging-promoting SASP factors granulocyte-macrophage colony-stimulating factor (GM-CSF), p = 0.05 (Fig. 6H) and interleukin-1α (IL-1α), p = 0.06 (Fig. 6I). In contrast, levels of granulocyte colony-stimulating factor (G-CSF), tumor necrosis factor receptor superfamily member 6 (TNFRSF6, also known as Fas or CD95), and vascular endothelial growth factor A (VEGF-A) proteins were reduced in aged p38β^-/-^ hearts compared to WT controls (Figs. 6F, 6L, and 6M), collectively indicating a reduced capacity for cardiac repair, including capillary repair, and increased vulnerability to cardiac stress during aging (21,22). Overall, these findings demonstrate that p38β deficiency reshapes the myocardial inflammatory environment in aged mice, promoting a cytokine and chemokine profile consistent with enhanced inflammaging and maladaptive myocardial remodeling.

**Fig. 6.**
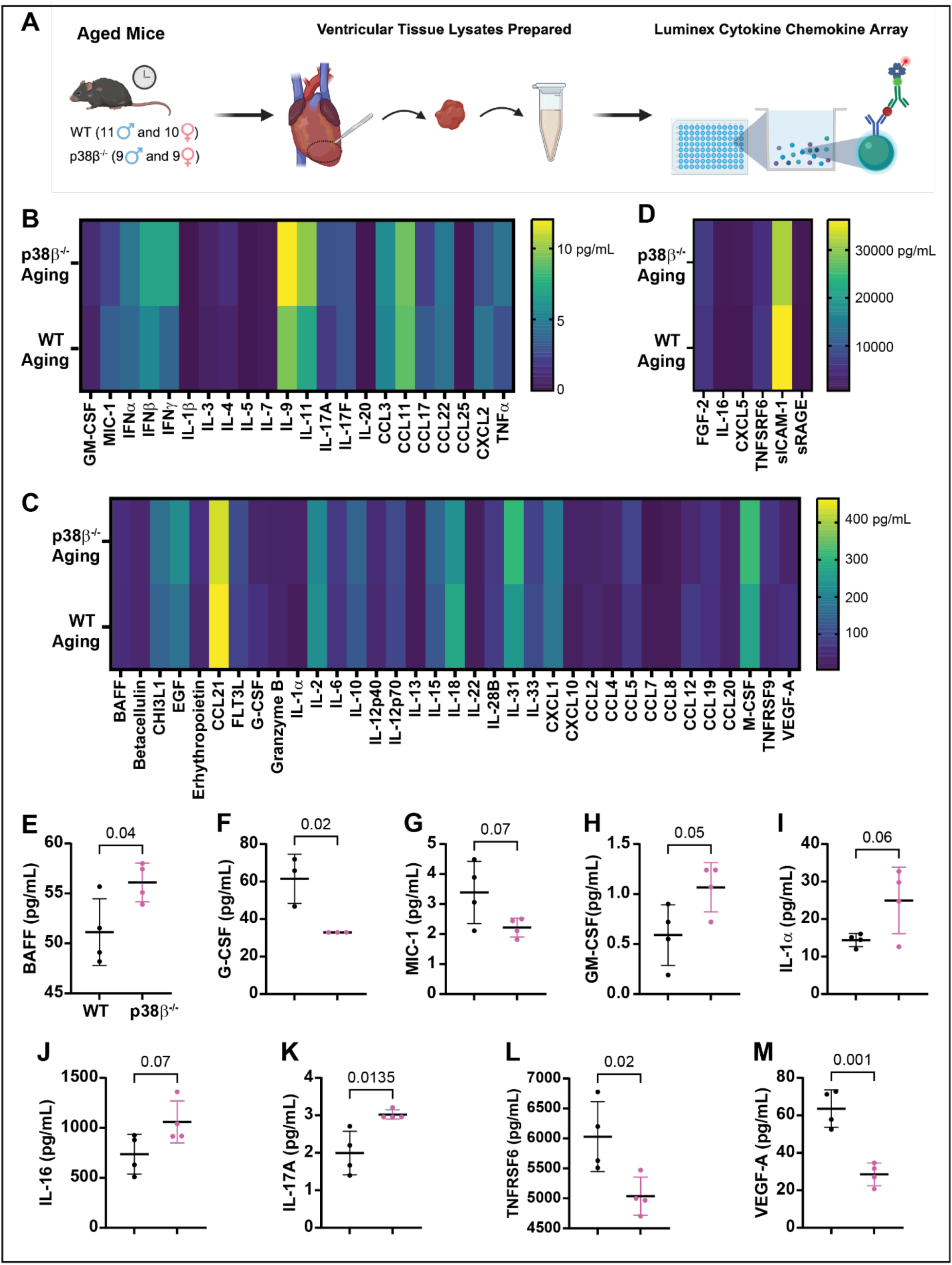
p38β deficiency promotes cardiac inflammation and inflammaging in aged mice. *A:* Schematic of the experimental protocol. *B – D*: Heatmap of chemokine and cytokine expression profiles in the hearts of aged (17-month-old) WT and p38β^-/-^ mice. Ventricular tissue lysates (total protein concentration 4 mg/ml) were used for mouse cytokine/chemokine array analysis. *E – M*: Molecules whose expression was significantly altered by p38β deficiency (p < 0.05) and those showing a trend toward higher expression (p = 0.05 to ≤ 0.07) are included. Data are presented as mean ± SD. An unpaired, two-tailed Student’s t-test was used to compare protein levels of cytokines and chemokines. Sample size: n = 4 mice per group. See also Supplemental Table 4.

### p38β deficiency promotes cardiac fibrosis in aged mice

Progressive fibrosis is a hallmark of cardiac aging and contributes to structural and mechanical remodeling of the myocardium (23). We assessed collagen deposition in the LV of aged WT and p38β^-/-^ mice using Picrosirius Red staining, as illustrated in Fig. 7A. Representative images show interstitial collagen deposition in the LVs of aged WT and p38β^-/-^ hearts (Fig. 7B). Fibrosis was quantified as the percentage of total tissue area occupied by collagen. Notably, aged p38β^-/-^ mice exhibited greater LV collagen deposition than their WT counterparts (Fig. 7C). These findings suggest that enhanced fibrotic remodeling may contribute to increased LV mass (Fig. 1E), LV wall thickness (Figs. 1F, 1K, and 1L), and myocardial stiffening in aged p38β^-/-^ mice compared with WT controls.

**Fig. 7.**
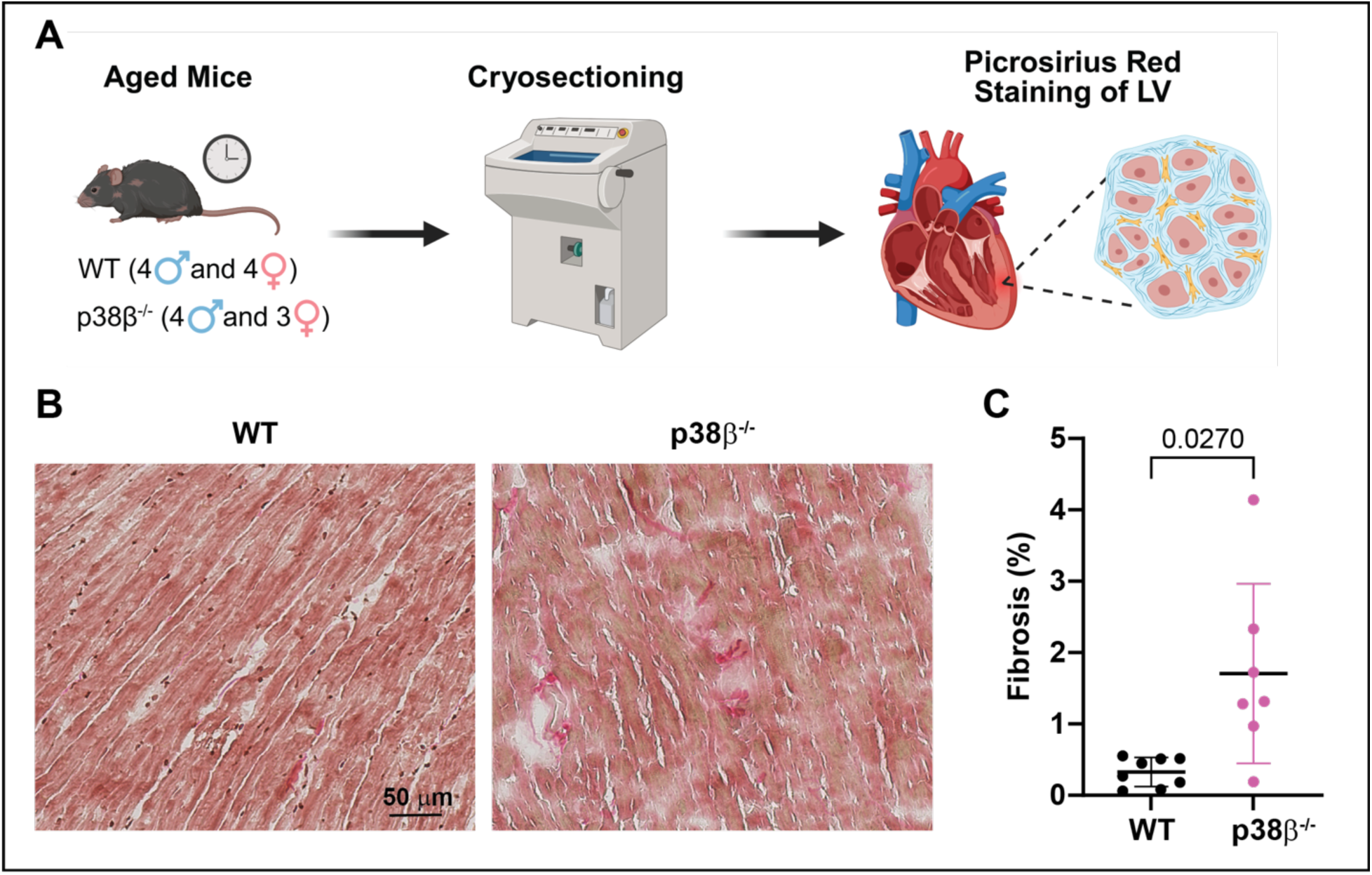
p38β deficiency promotes LV fibrosis in aged mice. *A:* Schematic of the experimental protocol. *B:* Representative images of LV tissue sections from aged (17-20-week-old) WT and p38β^-/-^ hearts stained with Picrosirius Red (collagen). *C:* Quantification of the percentage of LV area occupied by fibrosis. An unpaired, two-tailed Student’s t-test was used to assess statistical significance between the specified groups. Sample sizes: n = 8 in the WT group; n = 7 in the p38β^-/-^ group.

### p38β deficiency reprograms the cardiac transcriptome during aging

To investigate the molecular mechanisms underlying exacerbated structural, electrophysiological, and proinflammatory remodeling in aged p38β^-/-^ hearts, bulk RNA sequencing was performed on ventricular tissue from 17-month-old WT and p38β^-/-^ male mice, as illustrated in Fig. 8A. Principal component analysis revealed a clear separation between genotypes, with PC1 accounting for 51% of the total variance, indicating that p38β deletion is a major determinant of the aging cardiac transcriptome (Fig. 8B).

**Figure 8.**
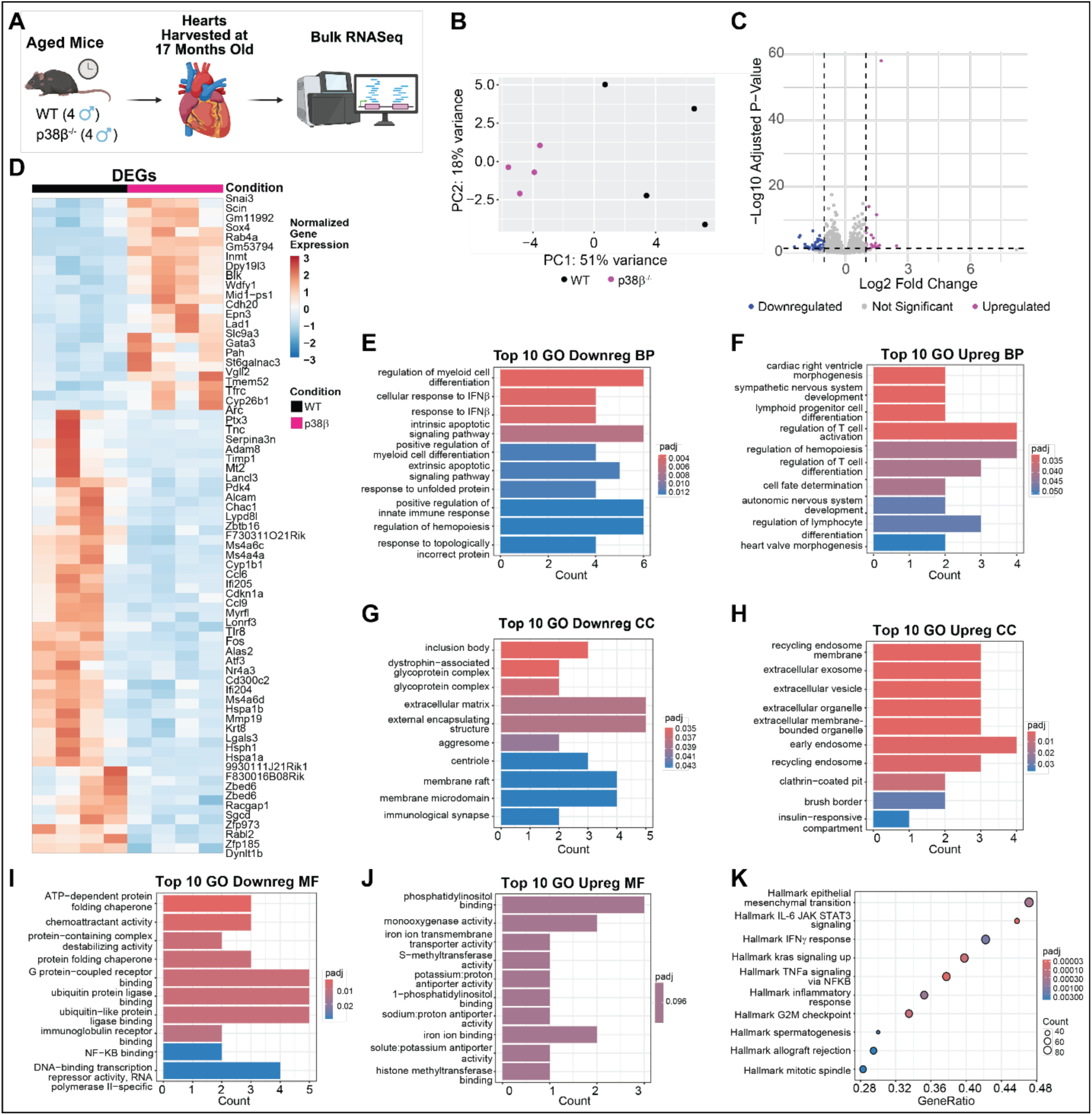
p38β deficiency regulates the cardiac transcriptome during aging. *A:* Schematic of the experimental protocol. *B:* Principal component analysis shows clear separation between genotypes. *C:* Volcano plot highlighting downregulated (blue) and upregulated (magenta) DEGs in p38β^-/-^ samples compared to WT controls. DEGs were identified using the DESeq2 package in R with a significance cutoff of padj < 0.1, and a log2 fold change > 1 for upregulated and < -1 for downregulated genes. *D:* Heatmap of normalized gene expression for DEGs, illustrating distinct transcriptional profiles between groups. *E – J:* Gene Ontology (GO) enrichment analysis of downregulated and upregulated genes in p38β^-/-^ hearts for Biological Process (BP), Cellular Component (CC), and Molecular Function (MF). The EnrichGO package in R was used to identify enriched GO pathways. *K:* Gene Set Enrichment Analysis (GSEA) of Hallmark pathways showing the top ten suppressed gene sets in aged p38β^-/-^ hearts. The ClusterProfiler package in R and the Hallmark Signatures Database (MSigDB, category H) were used for GSEA. An FDR q-value < 0.05 was considered significant for enrichment. Sample size: n = 4 per group. DEG: differentially expressed genes; padj: p-adjusted value; FDR: false discovery rate.

Differential expression analysis identified 68 differentially expressed genes (DEGs) in aged p38β^-/-^ hearts compared with their WT counterparts, of which 22 were upregulated and 46 were downregulated (Figs. 8C, D). Upregulated genes include structural and developmental genes such as *Snai3*, *Sox4*, *Cdh20*, *Vgll2*, and *Gata3*, suggesting aberrant activation of developmental pathways in aged p38β ^-/-^ hearts. Downregulated genes include immediate-early response genes, apoptosis regulators, and stress markers, such as *Fos*, *Atf3*, *Arc*, *Cdkn1a*, *Timp1*, and *Mt2*, as well as the key metabolic regulator *Pdk4* and protein-folding chaperones *Hspa1a*, *Hspa1b*, and *Hsph1*, indicating aberrant stress response, impaired cardiac energy metabolism, and impaired proteostasis (the balance of protein synthesis, folding, and degradation).

Gene Ontology Biological Process (GO BP) enrichment analysis of DEGs revealed that biological pathways related to interferon-β signaling (IFNβ), protein folding, proteostasis, response to unfolded or misfolded proteins, and innate immune responses, including regulation of myeloid cell differentiation, were significantly overrepresented among the downregulated genes (Fig. 8E). In contrast, biological pathways involved in cardiac morphogenesis, cell fate determination, hematopoiesis, regulation of T cell activation and differentiation, and autonomic and sympathetic nervous system development were significantly overrepresented among the upregulated genes (Fig. 8F), suggesting activation of developmental and neurohormonal remodeling programs.

Gene Ontology Cellular Component (GO CC) analysis showed that downregulated genes were enriched for GO CC terms related to extracellular matrix (ECM) remodeling and membrane-associated complexes, including the mechanoprotective dystrophin-associated glycoprotein complex (Fig. 8G), indicating changes in the myocardium’s structural organization and increased mechanical fragility of the p38β^-/-^ myocardium. Conversely, upregulated genes were enriched for GO CC terms associated with extracellular vesicles, exosomes, and endosome-related compartments (Fig. 8H), indicating altered membrane trafficking and intercellular signaling.

Gene Ontology Molecular Function (GO MF) analysis further showed that downregulated genes were enriched for GO MF terms related to protein folding, chaperone activity, ubiquitin ligase binding, and NFκB binding (Fig. 8I), consistent with impaired proteostasis. Loss of proteostasis is a recognized feature of age-related cardiovascular diseases, including atherosclerosis, atrial fibrillation, hypertrophic cardiomyopathy, and heart failure, in which an imbalance in protein synthesis, folding, and degradation contributes to cardiomyocyte stress (24). In contrast, upregulated genes were enriched for GO MF terms related to ion transport, including potassium/proton antiporter activity, sodium/proton exchange, phosphatidylinositol binding, iron ion transmembrane transporter activity, and monooxygenase activity (Fig. 8J), suggesting alterations in ion homeostasis and redox-related processes.

No KEGG pathways were significantly enriched. However, Gene Set Enrichment Analysis (GSEA) revealed suppression of Hallmark gene sets associated with epithelial-mesenchymal transition, IL6-JAK-STAT3 signaling, interferon-γ (IFNγ) response, kras signaling up, TNFα signaling via NFκB, inflammatory response, G2M checkpoint, spermatogenesis, allograft rejection, and mitotic spindle (Fig. 8K, Supplemental Tables 5 - 14).

Collectively, these findings suggest immune exhaustion, dysregulation of stress-response signaling, reduced regenerative capacity, electrical instability, proteotoxicity, and maladaptive tissue remodeling, including aberrant reactivation of an embryonic gene program, in aged p38β-deficient hearts compared with WT counterparts.

### p38β deficiency reduces trabecular bone volume in aged mice

CVD and osteoporosis are two highly prevalent age-related conditions increasingly recognized as interconnected (25). Clinical studies have shown that reduced bone mineral density is associated with adverse cardiac remodeling and correlated with heart failure risk in older adults (25). Given the pronounced cardiac remodeling observed in p38β^-/-^ mice, as well as the previously reported decreased bone mass in young (postnatal day 9 and 4-5-week-old) p38β^-/-^ mice (26), we next investigated whether loss of p38β affects bone integrity in aged mice. To assess skeletal remodeling, microCT was performed on aged (17-month-old) male WT, p38β^+/-,^ and p38β^-/-^ mice. Representative images of the left femur and zoomed-in axial views of cortical bone and distal metaphyseal trabecular bone for the specified genotypes are shown in Figs 9A and 9B. We found that the left femoral distal metaphyseal bone volume to total volume (BV/TV) was reduced in aged p38β^-/-^ mice and trended lower in aged p38β^+/-^ mice compared to their WT counterparts (Figs. 9B, 9C). No differences were observed in left or right femoral cortical bone thickness among the groups (Figs. 9D and 9E). These data indicate that p38β deficiency preferentially affects trabecular rather than cortical bone compartments in aged mice. Representative interparietal skull images from the specified groups are shown in Fig. 9F. Our analysis showed that interparietal bone BV/TV was significantly reduced in aged p38β^+/-^ mice compared to WT controls (Fig. 9H), suggesting decreased cranial bone mineralization, whereas skull volume was similar across all experimental groups (Fig. 9G). These findings are consistent with previously reported reductions in femoral trabecular BV/TV and calvarial mineralization in young p38β^-/-^ mice (26), and support a critical role for p38β in maintaining skeletal homeostasis during aging.

**Figure 9.**
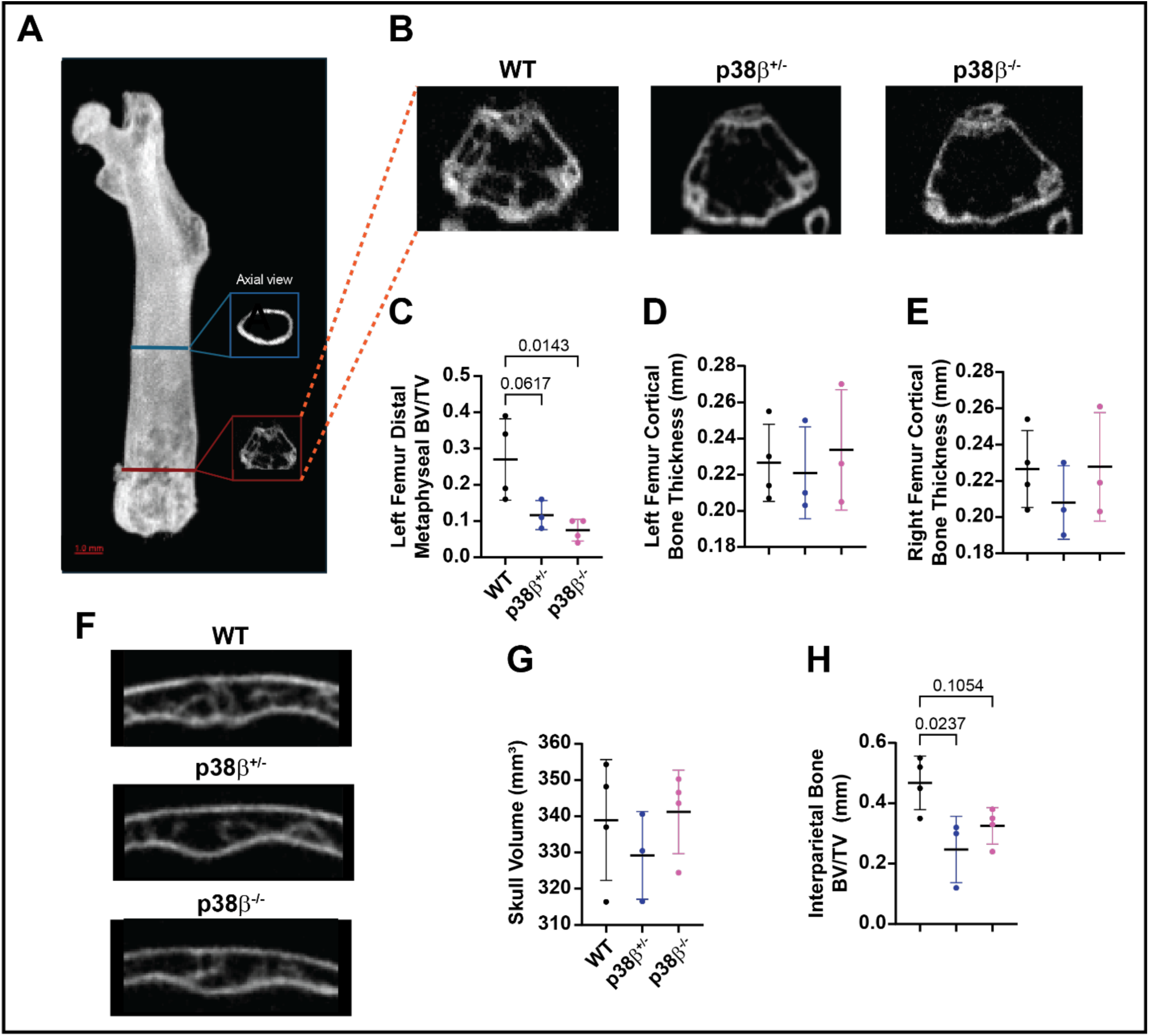
p38β deficiency reduces trabecular bone volume in aged mice. ***A:*** Representative microCT image of the left femur and zoomed-in axial views of cortical bone (blue box) and distal metaphyseal trabecular bone (red box) from aged mice. ***B:*** Representative microCT images of the distal metaphysis of the left femur from 17-month-old WT, p38β^+/-^, and p38β^-/-^ aged male mice. *C – E:* Quantification of left distal metaphyseal BV/TV **(C)**, left femur cortical bone thickness **(D)**, and right femur cortical bone thickness **(E)**. ***F:*** Representative microCT images of the interparietal skull bone from 17-month-old WT, p38β^+/-^, and p38β^-/-^ aged male mice. ***G, H:*** Quantification of skull volume **(G)** and interparietal bone BV/TV **(H)**. A one-way ANOVA with multiple comparisons was performed to compare across genotypes. Sample sizes: n = 4 in the WT group; n = 3 in the p38β^+/-^ group; n = 4 in the p38β^-/-^ group.

## DISCUSSION

Cardiac aging is characterized by progressive structural remodeling, electrophysiological dysfunction, and chronic low-grade inflammation, collectively increasing susceptibility to heart failure and arrhythmias. In this study, we identify p38β as a critical regulator of cardiac aging. p38β deficiency exacerbated cardiac electrophysiological and structural remodeling, including increased LV hypertrophy and fibrosis, impaired calcium handling, and heightened susceptibility to arrhythmias. Furthermore, p38β loss significantly altered the immune microenvironment in the aging heart (Fig. 10). Transcriptomic profiling revealed that p38β loss suppressed canonical innate immune and stress-response pathways, while promoting developmental, adaptive immune, and membrane signaling programs. Together, these findings demonstrate that p38β plays a cardioprotective role in the aging heart and that its loss drives maladaptive age-related remodeling and arrhythmogenesis.

**Figure 10.**
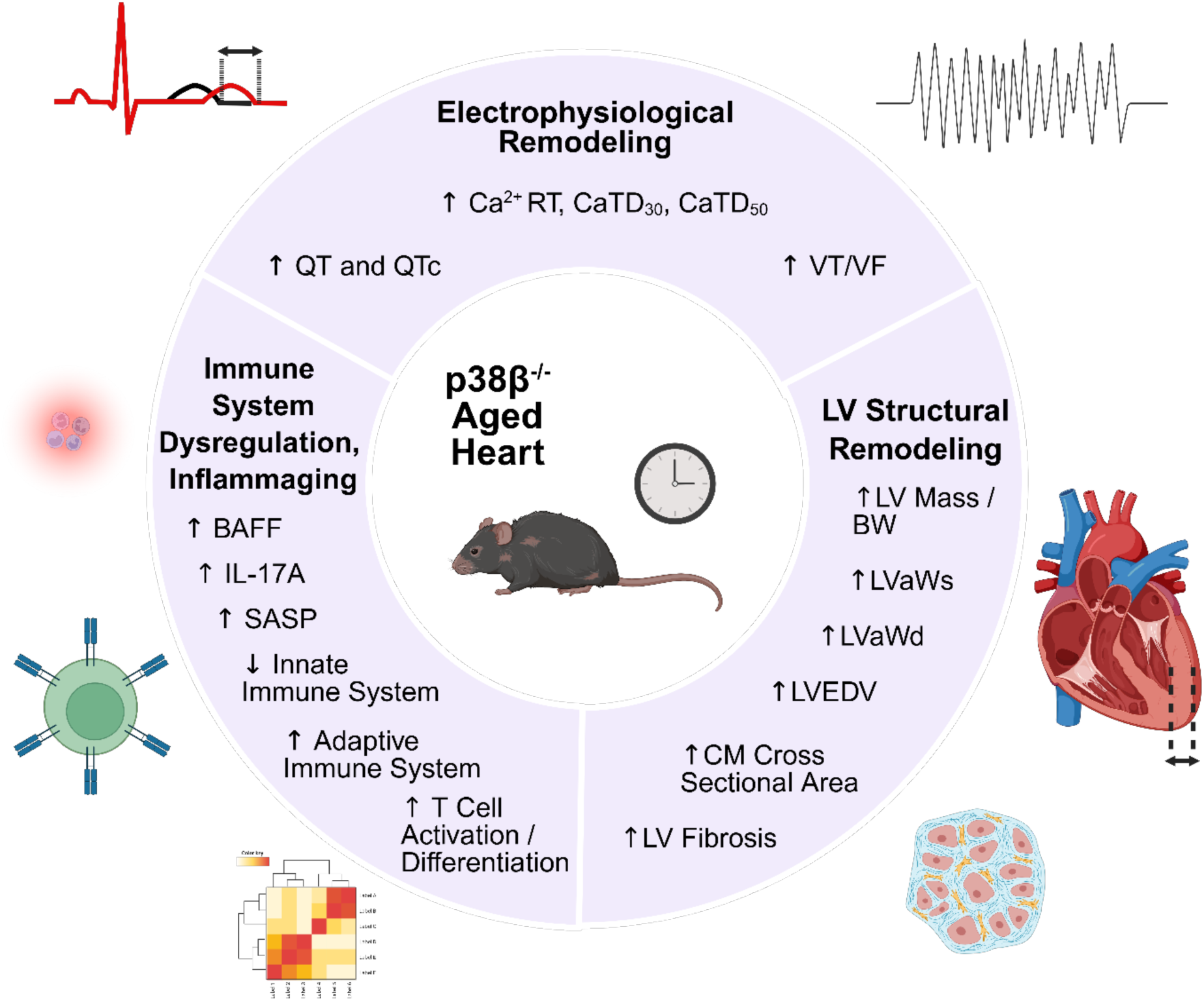
p38β deficiency accelerates cardiac aging and maladaptive remodeling. Loss of p38β promotes electrophysiological and LV structural remodeling, immune dysregulation and inflammaging, and global transcriptional reprogramming in aged mice.

Aging is associated with progressive LV structural remodeling. Even in healthy individuals, LV wall thickness and overall heart mass increase, indicating age-related cardiac hypertrophy (27–29). Importantly, these structural changes often occur without overt systolic dysfunction in the early stages of cardiac aging. Similar patterns are observed in mice, where LV remodeling begins around 12–15 months of age (30), corresponding to middle age (∼40–50 human years), while systolic dysfunction typically develops later, beginning around 18 months (∼60 human years) (31). Since C57BL/6 mice have a median lifespan of ∼29-30 months (∼85 human years) (32), this middle-aged window is a relevant period to study early age-related cardiac remodeling while minimizing confounding comorbidities of old to very old age.

In this study, we examined mice with an average age of approximately 16 months and observed worse pathological LV remodeling in aged p38β^-/-^ mice than in age-matched WT controls. Specifically, p38β deletion was associated with increases in LV mass, LV anterior wall thickness, and cardiomyocyte cross-sectional area, indicating enhanced cardiomyocyte hypertrophy. Mechanistically, transcriptomic analysis revealed upregulation of developmental and cell fate–associated pathways in aged p38β^⁻/⁻^ hearts. Among these, *Sox4*, a transcription factor that regulates tissue growth and differentiation, was increased, consistent with prior reports demonstrating its upregulation in hypertrophic hearts and the attenuation of cardiac hypertrophy following its knockdown (33).

Despite structural changes, overt LV systolic dysfunction was not observed. Instead, aged p38β^-/-^ mice showed increased SV, suggesting a compensatory response to ongoing structural remodeling. These findings align with previous research demonstrating that p38β deficiency worsens LV remodeling in models of cardiac stress (11–14). Notably, doxorubicin-induced cardiotoxicity has been described as a phenotype that mimics accelerated cardiac aging (8). In this regard, p38β deletion similarly promoted exaggerated LV wall thickening and reduced LV chamber diameter in female mice in a murine model of acute DIC (14).

Aging is associated with a progressive increase in ventricular arrhythmias, both in the presence and absence of structural heart disease (4). To avoid confounding by structural heart disease, we used a mouse model of natural aging to examine how loss of p38β affects cardiac electrophysiology during aging. Our data showed that p38β deficiency prolonged QT and QTc intervals in conscious aging mice, indicating impaired ventricular repolarization. QT prolongation is common in cardiac aging and is associated with Torsades de Pointes and sudden cardiac death (4,5,34). However, ex vivo optical mapping showed no differences in APD_30_, APD_50_, or APD_80_ across BCL, despite QT prolongation observed in vivo. This discrepancy suggests that in vivo QT prolongation is not due to inherent APD prolongation under isolated-heart conditions but rather results from systemic influences such as altered autonomic tone, β-adrenergic signaling, or inflammation, all of which were also identified as dysregulated at the transcriptomic level in aged p38β^-/-^ mice. Therefore, p38β modulates ventricular repolarization under neurohormonal stress.

Cardiac excitation-contraction coupling (ECC) is a highly coordinated process that couples transmembrane depolarization to mechanical contraction through tightly regulated Ca^2+^ cycling. Aging causes calcium mishandling by impairing calcium release, reuptake, and buffering, contributing to both contractile dysfunction and arrhythmogenesis (4,35). Notably, aged p38β-deficient hearts exhibited prolonged Ca^2+^ RT, indicating slowed Ca^2+^ release kinetics. This may reflect altered L-type Ca^2+^ channel (LTCC) activation or dysregulated sarcoplasmic reticulum Ca^2+^ release via ryanodine receptor 2 (RyR2). In the aged myocardium, RyR2 hyperactivity can promote triggered activity and premature ventricular complexes. Additionally, CaTD_30_ and CaTD_50_ were prolonged in aged p38β-deficient hearts compared with WT controls, indicating impaired calcium reuptake. This suggests potential alterations in SERCA2a function or its regulatory proteins. Reduced SERCA2a expression or alterations in its key regulator, phospholamban (PLB), such as reduced phosphorylation and increased PLB protein levels, are observed in the aging heart and may underlie the altered kinetics (35,36). Thus, p38β deficiency promotes a pro-arrhythmic electrophysiological substrate characterized by impaired calcium handling and reduced ventricular repolarization stability in the aged heart.

All aged p38β-deficient hearts developed VT/VF either spontaneously or during burst pacing, indicating increased susceptibility to arrhythmias. Although increased fibrosis was observed, optical activation maps and conduction velocity measurements showed largely preserved conduction, suggesting that arrhythmogenesis is not primarily due to widespread conduction slowing or fixed reentry. Instead, our findings support the notion of enhanced triggered activity. Aged p38β-deficient hearts exhibited a trend toward increased EAD incidence, consistent with ventricular repolarization instability. Impaired Ca^2+^ handling, including prolonged Ca^2+^ RT and delayed early reuptake, may elevate cytosolic Ca^2+^ and promote spontaneous SR release, which can generate inward current and facilitate afterdepolarizations (37). In addition, alterations in ion channel regulation and membrane trafficking may disrupt channel localization and destabilize V_m_, further contributing to electrical instability. Together, these findings suggest that p38β deficiency promotes a predominantly calcium- and ion-handling-driven focal arrhythmogenic substrate in the aged heart.

Epidemiological studies have identified inflammaging as a major risk factor for CVD and other age-related chronic diseases (17,38,39). Inflammaging is characterized by a chronic, low-grade proinflammatory state that serves as a marker of accelerated aging. As people age, the continuous production of proinflammatory cytokines, chemokines, growth factors, and extracellular vesicles alters the immune microenvironment, promoting persistent immune activation, cellular senescence, and extracellular matrix remodeling (40). Our findings show that loss of p38β amplifies multiple features consistent with an intensified cardiac inflammaging phenotype. Elevated IL-17A, along with trends toward higher GM-CSF and IL-1α, indicate heightened proinflammatory and SASP signaling in aged p38β-deficient hearts compared with WT counterparts. Circulating IL-17A increases with advancing age and in patients with heart failure (41), while GM-CSF and IL-1α are SASP factors that amplify inflammation (42). Lower VEGF-A levels in aging p38β-deficient hearts suggest impaired angiogenic and microvascular repair capacity, consistent with reports that VEGF insufficiency contributes to vascular aging and that restoring VEGF signaling ameliorates inflammaging (21,43). Increased expression of BAFF, a SASP component and a risk factor for coronary artery disease and acute myocardial infarction, further supports immune dysregulation in aged p38β-deficient hearts (19,44). Transcriptomic analysis further revealed pathological remodeling of the cardiac immune microenvironment, characterized by suppression of innate immune function alongside increased adaptive immune activation, lymphocyte recruitment, and hypersecretion of extracellular vesicles. During cardiac aging, T cell-mediated immune responses contribute to chronic low-grade inflammation through cytokine and chemokine production, enhanced SASP, and impaired clearance of senescent cells, thereby promoting a proinflammatory environment that, in turn, contributes to myocardial tissue damage and age-related cardiac dysfunction (45). Increased extracellular vesicle-associated pathways may further reflect enhanced SASP-mediated intercellular communication, thereby amplifying inflammatory signaling and adaptive immune recruitment during cardiac aging (40). Collectively, these coordinated alterations in inflammatory signaling suggest that the loss of p38β reshapes the cardiac immune microenvironment toward a more severe cardiac aging phenotype.

Osteoporosis is a common age-related disease characterized by progressive loss of bone mass due to an imbalance between bone resorption and bone formation (46). Increasing evidence suggests that osteoporosis and age-related CVD are linked through shared risk factors, chronic inflammation, and overlapping metabolic pathways that influence both the cardiovascular system and bone tissue (25). In this study, aged p38β-deficient mice exhibited reduced trabecular bone volume in the distal femur, indicating impaired maintenance of metabolically active bone with aging. In contrast, cranial bone changes were more modest, with reduced interparietal BV/TV but preserved overall skull volume. These findings are consistent with prior studies showing that young p38β^-/-^ mice display reduced femoral trabecular BV/TV and mild calvarial loss, and that p38β is required for late-stage osteoblast differentiation (26). The preferential remodeling of trabecular bone observed is consistent with age-related bone remodeling, in which trabecular bone is typically affected earlier and more severely than cortical bone (46). Likewise, long bones such as the femur are generally more vulnerable to age-related bone loss than craniofacial bones, which may explain the more pronounced bone loss in the femur in aged p38β-deficient mice (46). Collectively, these results further support p38β as a regulator of multiple organ systems during aging, contributing to both cardiovascular and skeletal homeostasis in aged mice.

This study examined mice aged 13.5 to 20 months. Mice in this age range have been reported to exhibit features of cardiac aging, including LV wall thickening, increased cardiomyocyte size, higher LV mass, increased oxidative stress, and worse outcomes with CVDs (47–49). Focusing on this age group allows assessment of age-related cardiac remodeling before severe aging occurs, a stage when survival bias and systemic comorbidities might skew results. However, the study did not include survival analysis, which prevents drawing conclusions about long-term outcomes and mortality. Future work should include survival analysis. Additionally, the use of a p38β germline KO model prevents determination of the effects of p38β loss in specific cell types. Since p38β is widely expressed, further studies using conditional, cell-type-specific transgenic approaches are needed to understand its roles in resident cardiac cells and infiltrating immune cells. Finally, future work should explore therapeutic strategies to enhance p38β expression or activity for cardioprotection, including investigation of recently identified small-molecule p38β activators such as DIPQUO (50).

### In summary

we demonstrate that p38β deficiency exacerbates cardiac aging, leading to increased hypertrophy, fibrosis, calcium mishandling, and heightened susceptibility to arrhythmias. Mechanistically, loss of p38β reprograms the cardiac transcriptome, suppressing innate immune response and proteostasis pathways while promoting adaptive immune activation, developmental remodeling, membrane trafficking, and ion transport signaling. These findings indicate that p38β regulates multiple processes central to cardiac aging, including immune homeostasis, structural remodeling, and electrophysiological stability. Together, our results reveal a previously unrecognized cardioprotective role for p38β in maintaining cardiac function during aging. Importantly, these findings suggest that non-selective inhibition of p38 MAPKs may disrupt the beneficial p38β-dependent signaling and exacerbate age-related cardiac dysfunction, highlighting the need for isoform-specific therapeutic strategies.

## NONSTANDARD ABBREVIATIONS AND ACRONYMS

APD_30_: Action Potential Duration at 30% Repolarization
APD_50_: Action Potential Duration at 50% Repolarization
APD_80_: Action Potential Duration at 80% Repolarization
AR_CV_: Anisotropic Conduction Velocity
BAFF: B-cell Activating Factor
BP: Biological Pathway
CaT: Calcium Transient
CaTD_30_: Calcium Transient Duration at 30% Reuptake
CaTD_50_: Calcium Transient Duration at 50% Reuptake
CaTD_80_: Calcium Transient Duration at 80% Reuptake
Ca^2+^ RT: Calcium Rise Time
Ca^2+^τ: Calcium Decay
CC: Cellular Component
CO: Cardiac Output
CVD: Cardiovascular Disease
CV_L_: Longitudinal Conduction Velocity
CV_T_: Transverse Conduction Velocity
DEG: Differentially Expressed Genes
ECC: Excitation-Contraction Coupling
ECG: Electrocardiogram
EF: Ejection Fraction
G-CSF: Granulocyte Colony-Stimulating Factor
GM-CSF: Granulocyte-Macrophage Colony-Stimulating Factors
GSEA: Gene Set Enrichment Analysis
IFNα: Interferon α
IFNβ: Interferon β
IFNγ: Interferon γ
IL-1α: Interleukin-1α
IL-17A: Interleukin-17A
KO: Knockout
LTCC: L-type Calcium Channel
LV: Left Ventricle
LVaWd: Left Ventricular Anterior Wall during Diastole
LVaWs: Left Ventricular Anterior Wall during Systole
LVEDD: Left Ventricular End-Diastolic Diameter
LVESD: Left Ventricular End-Systolic Diameter
LVEDV: Left Ventricular End-Diastolic Volume
LVESV: Left Ventricular End-Systolic Volume
LVpWd: Left Ventricular Posterior Wall during Diastole
LVpWs: Left Ventricular Posterior Wall during Systole
MAPK: Mitogen-Activated Protein Kinase
MF: Molecular Function
OAP: Optical Action Potential
RyR2: Ryanodine Receptor 2
SASP: Senescence-Associated Secretory Phenotype
TNFRSF6: Tumor Necrosis Factor Receptor Superfamily Member 6
VEGF-A: Vascular Endothelial Growth Factor A
VF: Ventricular Fibrillation
V_m_: Transmembrane Potential
V_m_ RT: Transmembrane Potential Rise Time
VT: Ventricular Tachycardia
WT: Wild-type

## Data Availability

Data will be made available upon reasonable request.

## ACKNOWLEDGEMENTS

We thank Dr. Paloma Amaral for her excellent technical assistance.

## AUTHOR CONTRIBUTIONS

Conceived and designed experiments – KAT, IRE, TE; performed experiments – KAT, BS, SG, AA; analyzed data – KAT, BS, SG, AA TE; interpreted results of experiments – KAT, IRE, TE; prepared figures – KAT, TE; drafted manuscript – KAT, SG, TE; edited and revised manuscript – KAT, TE, IRE; all the authors approved the final version of the manuscript.

## Supplementary Information

**Supplemental Table 1.** Summary of cardiac functional data in aged WT and p38β^-/-^mice. Parameters of cardiac structural, mechanical, and electrical functions measured by echocardiography and electrocardiography in aged WT and p38β^-/-^ mice are summarized. The groups were age- and sex-matched, with a mean age of approximately 16 months. Data are presented as mean ± SD. * *P* < 0.05, WT versus p38β^-/-^ groups. Sample sizes: n = 21 in the WT group; n = 18 in the p38β^-/-^ group. The data in this table are related to Figures 1 and 2.

| | WT | p38β<sup>-/-</sup> |
| --- | --- | --- |
| **Echocardiography** |  |  |
| EF, % | 30.33 ± 7.00 | 31.89 ± 4.75 |
| FS, % | 57.50 ± 10.39 | 59.71 ± 7.03 |
| SV, μL | 44.02 ± 11.52 | 51.89 ± 6.94* |
| CO, mL/min | 17.84 ± 4.76 | 18.68 ± 3.91 |
| HR (bpm) | 411.38 ± 65.77 | 359.35 ± 60.36 |
| LVaWs, mm | 1.39 ± 0.27 | 1.67 ± 0.27* |
| LVaWd, mm | 0.97 ± 0.20 | 1.20 ± 0.23* |
| LVpWs, mm | 1.30 ± 0.20 | 1.37 ± 0.24 |
| LVpWd, mm | 0.95 ± 0.22 | 0.98 ± 0.16 |
| LVEDS, mm | 2.89 ± 0.46 | 3.06 ± 0.55 |
| LVEDD, mm | 4.15 ± 0.41 | 4.26 ± 0.45 |
| LVESV, mL | 33.24 ± 13.30 | 36.04 ± 10.83 |
| LVEDV, mL | 77.28 ± 18.23 | 87.84 ± 15.21 |
| **Electrocardiography** |  |  |
| P, ms | 16.44 ± 2.44 | 16.87 ± 1.57 |
| PR, ms | 42.95 ± 3.45 | 41.82 ± 4.15 |
| QRS, ms | 10.34 ± 2.50 | 10.07 ± 1.69 |
| QT, ms | 30.37 ± 5.49 | 37.63 ± 3.05* |
| RR, ms | 85.50 ± 4.52 | 85.78 ± 2.74 |
| QTc, ms | 32.87 ± 5.97 | 40.63 ± 3.14* |

**Supplemental Table 2.** Summary of cardiac functional data in aged WT male and female mice. Parameters of cardiac structural, mechanical, and electrical functions measured by echocardiography and electrocardiography in aged WT male and female mice are summarized. Data are presented as mean ± SD. * *P* < 0.05, male versus female. Sample sizes: n = 11 in the WT male group; n = 10 in the WT female group. The data in this table are related to Figures 1 and 2.

|  | WT Males | WT Females |
| --- | --- | --- |
| **Echocardiography** |  |  |
| EF, % | 57.18 ± 11.17 | 57.84 ± 10.04 |
| FS, % | 30.21 ± 7.46 | 30.47 ± 6.87 |
| SV, μL | 47.76 ± 12.49 | 39.91 ± 9.26 |
| CO, mL/min | 19.57 ± 5.19 | 15.94 ± 3.57 |
| LVaWs, mm | 1.42 ± 0.22 | 1.36 ± 0.32 |
| LVaWd, mm | 0.98 ± 0.24 | 0.96 ± 0.17 |
| LVpWs, mm | 1.34 ± 0.22 | 1.26 ± 0.17 |
| LVpWd, mm | 0.97 ± 0.23 | 0.92 ± 0.22* |
| LVEDS, mm | 2.98 ± 0.38 | 2.79 ± 0.54 |
| LVEDD, mm | 4.28 ± 0.30 | 4.00 ± 0.48 |
| LVESV, mL | 35.67 ± 10.77 | 30.56 ± 15.78 |
| LVEDV, mL | 0.98 ± 0.24 | 0.96 ± 0.17 |
| **Electrocardiography** |  |  |
| P, ms | 16.13 ± 2.08 | 16.78 ± 2.87 |
| PR, ms | 41.60 ± 3.67 | 44.44 ± 2.61 |
| QRS, ms | 10.54 ± 2.67 | 10.12 ± 2.43 |
| QT, ms | 30.21 ± 4.85 | 30.56 ± 4.85 |
| RR, ms | 83.84 ± 3.17 | 87.32 ± 5.22 |
| QTc, ms | 33.02 ± 5.49 | 32.71 ± 6.75 |

**Supplemental Table 3.** Summary of cardiac functional data in aged p38β^-/-^ male and female mice. Parameters of cardiac structural, mechanical, and electrical functions measured by echocardiography and electrocardiography in aged p38β^-/-^ male and female mice are summarized. Data are presented as mean ± SD. * *P* < 0.05, p38β^-/-^ male versus p38β^-/-^ female mice. Sample size: n = 9 in each group. The data in this table are related to Figures 1 and 2.

| | p38β<sup>-/-</sup> Males | p38β<sup>-/-</sup> Females |
| --- | --- | --- |
| **Echocardiography** |  |  |
| EF, % | 59.97 ± 7.38 | 58.80 ± 6.90 |
| FS, % | 32.02 ± 5.14 | 31.20 ± 4.52 |
| SV, μL | 53.59 ± 6.06 | 50.55 ± 7.47 |
| CO, mL/min | 20.43 ± 3.49 | 17.39 ± 3.90 |
| LVaWs, mm | 1.67 ± 0.27 | 1.62 ± 0.33 |
| LVaWd, mm | 1.26 ± 0.26 | 1.09 ± 0.21 |
| LVpWs, mm | 1.45 ± 0.28 | 1.28 ± 0.18 |
| LVpWd, mm | 1.06 ± 0.17 | 0.88 ± 0.11 |
| LVEDS, mm | 3.16 ± 0.62 | 3.00 ± 0.46 |
| LVEDD, mm | 4.23 ± 0.54 | 4.33 ± 0.36 |
| LVESV, mL | 36.76 ± 10.92 | 36.58 ± 11.42 |
| LVEDV, mL | 90.34 ± 13.69 | 86.99 ± 17.18 |
| **Electrocardiography** |  |  |
| P, ms | 16.93 ± 1.94 | 16.81 ± 1.21 |
| PR, ms | 42.11 ± 3.99 | 41.53 ± 4.53 |
| QRS, ms | 10.19 ± 1.55 | 9.95 ± 1.91 |
| QT, ms | 36.53 ± 3.28 | 38.74 ± 2.51 |
| RR, ms | 84.61 ± 2.85 | 86.95 ± 2.18 |
| QTc, ms | 39.70 ± 3.23 | 41.56 ± 2.92 |

**Supplemental Table 4.** p38β deficiency modulates cardiac chemokine and cytokine expression profiles in aged mice. Protein levels of the indicated inflammatory mediators in the hearts of 17-month-old WT and p38β^-/-^ male mice. Total ventricular tissue lysates were used for cytokine and chemokine array analysis. Data are presented as mean ± SD. Unpaired two-tailed Student’s t-tests were used to compare inflammatory mediator levels between groups. * *P* < 0.05 (shaded in green) denotes statistical significance. *P* < 0.1 suggests a trend (shaded in blue). Sample size: n = 4 mice per group. GF: growth factor. The data in this table are related to Figure 6.

| Chemokine, Cytokine, or GF | Concentration (pg/mL) | | p38β<sup>-/-</sup> Aged / WT Aged | p value |
| --- | --- | --- | --- | --- |
| | WT Aged | p38β<sup>-/-</sup> Aged | | |
| **BAFF** | 51.13 ± 3.35 | 56.11 ± 1.94 | 1.10 | 0.04 |
| **Betacellulin** | 40.99 ± 7.29 | 37.80 ± 3.16 | 0.92 | 0.45 |
| **CCL25** | 0.00 | 0.00 | NA | NA |
| **CHI3L1** | 162.54 ± 63.38 | 161.48 ± 21.86 | 0.99 | 0.98 |
| **CXCL16** | 310.08 ± 136.34 | 424.90 ± 65.18 | 1.37 | 0.18 |
| **EGF** | 180.08 ± 42.01 | 225.26 ± 14.78 | 1.25 | 0.09 |
| **CCL11** | 9.27 ± 1.01 | 9.23 ± 0.35 | 1.00 | 0.95 |
| **Erythropoietin** | 42.42 ± 15.82 | 57.92 ± 9.65 | 1.37 | 0.15 |
| **CCL21** | 481.54 ± 95.29 | 462.73 ± 120.82 | 0.96 | 0.82 |
| **FGF-2** | 6176.43 ± 997.37 | 6912.62 ± 1338.91 | 1.12 | 0.41 |
| **FLT3L** | 100.06 ± 23.97 | 108.83 ± 15.85 | 1.09 | 0.56 |
| **CX3CL1** | 42.54 ± 7.78 | 45.59 ± 3.29 | 1.07 | 0.50 |
| **G-CSF** | 61.51 ± 13.12 | 32.88 ± 0.00 | 0.53 | 0.02 |
| **MIC-1** | 3.39 ± 1.03 | 2.22 ± 0.32 | 0.65 | 0.07 |
| **GM-CSF** | 0.59 ± 0.30 | 1.07 ± 0.25 | 1.81 | 0.05 |
| **Granzyme B** | 31.89 ± 13.13 | 29.64 ± 9.76 | 0.93 | 0.79 |
| **IFNα** | 4.19 ± 1.14 | 4.68 ± 1.06 | 1.12 | 0.55 |
| **IFNβ** | 5.37 ± 2.02 | 6.87 ± 0.50 | 1.28 | 0.20 |
| **IFNγ** | 4.37 ± 2.40 | 6.85 ± 3.56 | 1.57 | 0.29 |
| **IL-1α** | 14.40 ± 1.74 | 24.95 ± 8.87 | 1.73 | 0.06 |
| **IL-1β** | 0.00 | 0.00 | NA | NA |
| **IL-2** | 223.85 ± 25.22 | 219.89 ± 25.70 | 0.98 | 0.83 |
| **IL-3** | 0.52 ± 0.12 | 0.68 ± 0.14 | 1.31 | 0.14 |
| **IL-4** | 0.68 ± 0.47 | 1.26 ± 0.37 | 1.84 | 0.10 |
| **IL-5** | 0.00 | 0.00 | NA | NA |
| **IL-6** | 69.58 ± 11.01 | 81.01 ± 11.42 | 1.16 | 0.20 |
| **IL-7** | 0.20 ± 0.03 | 0.19 ± 0.05 | 0.93 | 0.59 |
| **IL-9** | 11.29 ± 5.12 | 11.39 ± 3.21 | 1.01 | 0.97 |
| **IL-10** | 115.06 ± 33.00 | 149.85 ± 28.00 | 1.30 | 0.16 |
| **IL-11** | 7.57 ± 3.70 | 10.21 ± 1.16 | 1.35 | 0.22 |
| **IL-12p40** | 42.77 ± 10.58 | 50.89 ± 6.30 | 1.19 | 0.24 |
| **IL-12p70** | 98.15 ± 27.36 | 108.11 ± 17.70 | 1.10 | 0.56 |
| **IL-13** | 13.20 ± 4.51 | 16.94 ± 3.63 | 1.28 | 0.24 |
| **IL-15** | 130.05 ± 29.42 | 151.44 ± 34.25 | 1.16 | 0.38 |
| **IL-16** | 736.48 ± 199.04 | 1058.84 ± 209.95 | 1.44 | 0.07 |
| **IL-17A** | 2.00 ± 0.58 | 3.02 ± 0.13 | 1.52 | 0.01 |
| **IL-17F** | 3.11 ± 0.25 | 3.23 ± 0.61 | 1.04 | 0.71 |
| **IL-18** | 267.83 ± 22.81 | 245.21 ± 51.53 | 0.92 | 0.45 |
| **IL-20** | 0.00 | 0.00 | NA | NA |
| **IL-22** | 16.26 ± 3.56 | 16.80 ± 2.18 | 1.03 | 0.80 |
| **IL-28B** | 83.39 ± 3.52 | 82.07 ± 10.91 | 0.98 | 0.83 |
| **IL-31** | 238.25 ± 98.71 | 336.56 ± 93.53 | 1.41 | 0.20 |
| **IL-33** | 99.29 ± 12.89 | 89.27 ± 4.40 | 0.90 | 0.19 |
| **CXCL10** | 19.53 ± 8.90 | 27.43 ± 5.47 | 1.40 | 0.18 |
| **CXCL1** | 180.34 ± 15.05 | 198.19 ± 36.02 | 1.10 | 0.40 |
| **LIF** | 0.78 ± 1.00 | 1.62 ± 0.78 | 2.09 | 0.23 |
| **CXCL5** | 721.75 ± 369.67 | 1121.08 ± 615.41 | 1.55 | 0.31 |
| **M-CSF** | 268.54 ± 71.03 | 340.89 ± 64.28 | 1.27 | 0.18 |
| **CCL2** | 51.64 ± 26.50 | 26.97 ± 10.96 | 0.52 | 0.14 |
| **CCL8** | 9.97 ± 2.77 | 20.95 ± 20.26 | 2.10 | 0.32 |
| **CCL7** | 17.32 ± 11.76 | 5.44 ± 1.17 | 0.31 | 0.09 |
| **CCL12** | 97.23 ± 53.69 | 52.89 ± 25.51 | 0.54 | 0.19 |
| **CCL22** | 4.59 ± 1.05 | 4.94 ± 0.76 | 1.08 | 0.60 |
| **CXCL9** | 92.63 ± 39.24 | 148.22 ± 38.98 | 1.60 | 0.09 |
| **CCL3** | 5.75 ± 1.22 | 5.66 ± 0.71 | 0.98 | 0.90 |
| **CCL4** | 25.64 ± 6.18 | 37.93 ± 13.27 | 1.48 | 0.14 |
| **CXCL2** | 4.59 ± 2.77 | 3.24 ± 0.34 | 0.71 | 0.11 |
| **CCL20** | 28.61 ± 17.29 | 29.29 ± 12.66 | 1.02 | 0.83 |
| **CCL19** | 78.61 ± 24.41 | 75.88 ± 41.61 | 0.97 | 0.59 |
| **CCL5** | 91.94 ± 17.13 | 73.28 ± 19.67 | 0.80 | 0.15 |
| **TNFRSF9** | 71.83 ± 26.64 | 77.97 ± 20.57 | 1.09 | 0.11 |
| **TNFSRF6** | 5509.01 ± 82.60 | 4968.11 | 0.90 | 0.02 |
| **sFasL** | 33.21 ± 5.65 | 44.22 ± 8.31 | 1.33 | 0.33 |
| **sICAM-1** | 35478.26 ±<br>4303.41 | 30367.28 ±<br>2778.24 | 0.86 | 0.09 |
| **sRAGE** | 620.82 ± 383.54 | 921.66 ± 202.38 | 1.48 | 0.21 |
| **CCL17** | 3.26 ± 0.89 | 2.20 ± 0.68 | 0.67 | 0.10 |
| **TNFα** | 2.96 ± 1.61 | 4.07 ± 1.00 | 1.37 | 0.29 |
| **VEGF-A** | 63.58 ± 9.98 | 28.56 ± 6.04 | 0.45 | 0.001 |

**Supplemental Table 5.** Suppression of the Hallmark Epithelial Mesenchymal Transition gene set in aged p38β^-/-^ hearts. The Hallmark Epithelial Mesenchymal Transition gene set was identified by GSEA as one of the top 10 suppressed gene sets in the hearts of aged p38β^-/-^ male mice compared with WT counterparts. The data in this table are related to Figure 8J.

| Gene Symbol | Gene Name |
| --- | --- |
| Rgs4 | Regulator of G Protein Signaling 4 |
| Itgb1 | Integrin Subunit Beta 1 |
| Emp3 | Epithelial Membrane Protein 3 |
| Ccn2 | Cellular Communication Network Factor 2 (CTGF) |
| Lamc1 | Laminin Subunit Gamma 1 |
| Fn1 | Fibronectin 1 |
| Slit2 | Slit Guidance Ligand 2 |
| Loxl2 | Lysyl Oxidase Like 2 |
| Tpm4 | Tropomyosin 4 |
| Fbn1 | Fibrillin 1 |
| Serpine2 | Serpin Family E Member 2 |
| Cxcl1 | C-X-C Motif Chemokine Ligand 1 |
| Fbln2 | Fibulin 2 |
| Ntm | Neurotrimin |
| Pmp22 | Peripheral Myelin Protein 22 |
| Pvr | Poliovirus Receptor |
| Col6a2 | Collagen Type VI Alpha 2 Chain |
| Lrp1 | LDL Receptor Related Protein 1 |
| Col4a1 | Collagen Type IV Alpha 1 Chain |
| Col8a2 | Collagen Type VIII Alpha 2 Chain |
| Sparc | Secreted Protein Acidic And Cysteine Rich |
| Dst | Dystonin |
| Cdh11 | Cadherin 11 |
| Pcolce | Procollagen C-Endopeptidase Enhancer |
| Gpx7 | Glutathione Peroxidase 7 |
| Loxl1 | Lysyl Oxidase Like 1 |
| Lum | Lumican |
| Plod1 | Procollagen-Lysine,2-Oxoglutarate 5-Dioxygenase 1 |
| Col5a1 | Collagen Type V Alpha 1 Chain |
| Matn2 | Matrilin 2 |
| Acta2 | Actin Alpha 2, Smooth Muscle |
| Basp1 | Brain Abundant Membrane Attached Signal Protein 1 |
| Mgp | Matrix Gla Protein |
| Dab2 | DAB2, Clathrin Adaptor Protein |
| Plod3 | Procollagen-Lysine,2-Oxoglutarate 5-Dioxygenase 3 |
| Fmod | Fibromodulin |
| Adam12 | ADAM Metallopeptidase Domain 12 |
| Lama2 | Laminin Subunit Alpha 2 |
| Mmp2 | Matrix Metallopeptidase 2 |
| Abi3bp | ABI Family Member 3 Binding Protein |
| Timp3 | TIMP Metallopeptidase Inhibitor 3 |
| Vcam1 | Vascular Cell Adhesion Molecule 1 |
| Col16a1 | Collagen Type XVI Alpha 1 Chain |
| Itga2 | Integrin Subunit Alpha 2 |
| Itgav | Integrin Subunit Alpha V |
| Capg | Capping Actin Protein Gelsolin Like |
| Ecm1 | Extracellular Matrix Protein 1 |
| Fstl1 | Follistatin Like 1 |
| Vim | Vimentin |
| Aplp1 | Amyloid Beta Precursor Like Protein 1 |
| Bgn | Biglycan |
| Lamc2 | Laminin Subunit Gamma 2 |
| Fstl3 | Follistatin Like 3 |
| Col5a2 | Collagen Type V Alpha 2 Chain |
| Inhba | Inhibin Subunit Beta A |
| Col1a1 | Collagen Type I Alpha 1 Chain |
| Tnfrsf12a | TNF Receptor Superfamily Member 12A |
| Fbln1 | Fibulin 1 |
| Fas | Fas Cell Surface Death Receptor |
| Col1a2 | Collagen Type I Alpha 2 Chain |
| Col3a1 | Collagen Type III Alpha 1 Chain |
| Col12a1 | Collagen Type XII Alpha 1 Chain |
| Postn | Periostin |
| Itgb3 | Integrin Subunit Beta 3 |
| Cd44 | CD44 Molecule |
| Col6a3 | Collagen Type VI Alpha 3 Chain |
| Scg2 | Secretogranin II |
| Serpinh1 | Serpin Family H Member 1 (HSP47) |
| Edil3 | EGF Like Repeats And Discoidin Domains 3 |
| Mfap5 | Microfibril Associated Protein 5 |
| Lox | Lysyl Oxidase |
| Glipr1 | GLI Pathogenesis Related 1 |
| Plaur | Plasminogen Activator, Urokinase Receptor |
| Serpine1 | Serpin Family E Member 1 (PAI-1) |
| Mmp3 | Matrix Metallopeptidase 3 |
| Sfrp4 | Secreted Frizzled Related Protein 4 |
| Crlf1 | Cytokine Receptor Like Factor 1 |
| Cdh6 | Cadherin 6 |
| Vcan | Versican |
| Thbs1 | Thrombospondin 1 |
| Cthrc1 | Collagen Triple Helix Repeat Containing 1 |
| Sgcd | Sarcoglycan Delta |
| Cxcl2 | C-X-C Motif Chemokine Ligand 2 |
| Cxcl3 | C-X-C Motif Chemokine Ligand 3 |
| Tnfrsf11b | TNF Receptor Superfamily Member 11B (Osteoprotegerin) |
| Ptx3 | Pentraxin 3 |
| Timp1 | TIMP Metallopeptidase Inhibitor 1 |
| Il6 | Interleukin 6 |
| Tnc | Tenascin C |
| Spock1 | SPARC/Osteonectin, Cwcv And Kazal Like Domains Proteoglycan 1 |
| Spp1 | Secreted Phosphoprotein 1 (Osteopontin) |

**Supplemental Table 6.** Suppression of the Hallmark IL-6 JAK STAT3 Signaling gene set in aged p38β^-/-^ hearts. The Hallmark IL-6 JAK STAT3 Signaling gene set was identified by GSEA as one of the top 10 suppressed gene sets in the hearts of aged p38β^-/-^ male mice compared with WT counterparts. The data in this table are related to Figure 8J.

| Gene Symbol | Gene Name |
| --- | --- |
| Cd9 | CD9 Molecule |
| Map3k8 | Mitogen-Activated Protein Kinase Kinase Kinase 8 (TPL2) |
| Il17rb | Interleukin 17 Receptor B |
| Myd88 | MYD88 Innate Immune Signal Transduction Adaptor |
| Ccr1 | C-C Motif Chemokine Receptor 1 |
| Cbl | Cbl Proto-Oncogene |
| Stat3 | Signal Transducer and Activator of Transcription 3 |
| Il1b | Interleukin 1 Beta |
| Il4ra | Interleukin 4 Receptor Subunit Alpha |
| Csf3r | Colony Stimulating Factor 3 Receptor |
| Cntfr | Ciliary Neurotrophic Factor Receptor |
| Tnfrsf12a | TNF Receptor Superfamily Member 12A (Fn14) |
| Il2ra | Interleukin 2 Receptor Subunit Alpha (CD25) |
| Fas | Fas Cell Surface Death Receptor |
| Il12rb1 | Interleukin 12 Receptor Subunit Beta 1 |
| Il17ra | Interleukin 17 Receptor A |
| Pik3r5 | Phosphoinositide-3-Kinase Regulatory Subunit 5 |
| Itgb3 | Integrin Subunit Beta 3 |
| Cd44 | CD44 Molecule |
| Pdgfc | Platelet Derived Growth Factor C |
| Tnf | Tumor Necrosis Factor |
| Osmr | Oncostatin M Receptor |
| Cd14 | CD14 Molecule |
| Hmox1 | Heme Oxygenase 1 |
| Cxcl10 | C-X-C Motif Chemokine Ligand 10 |
| Csf2rb2 | Colony Stimulating Factor 2 Receptor Beta 2 |
| Itga4 | Integrin Subunit Alpha 4 |
| Cxcl13 | C-X-C Motif Chemokine Ligand 13 |
| Ccl7 | C-C Motif Chemokine Ligand 7 |
| Il18r1 | Interleukin 18 Receptor 1 |
| Cxcl2 | C-X-C Motif Chemokine Ligand 2 |
| Socs3 | Suppressor Of Cytokine Signaling 3 |
| Cxcl3 | C-X-C Motif Chemokine Ligand 3 |
| Csf2 | Colony Stimulating Factor 2 (GM-CSF) |
| Il6 | Interleukin 6 |
| A2m | Alpha-2-Macroglobulin |
| Il1r2 | Interleukin 1 Receptor Type 2 |
| Inhbe | Inhibin Subunit Beta E |

**Supplemental Table 7.**
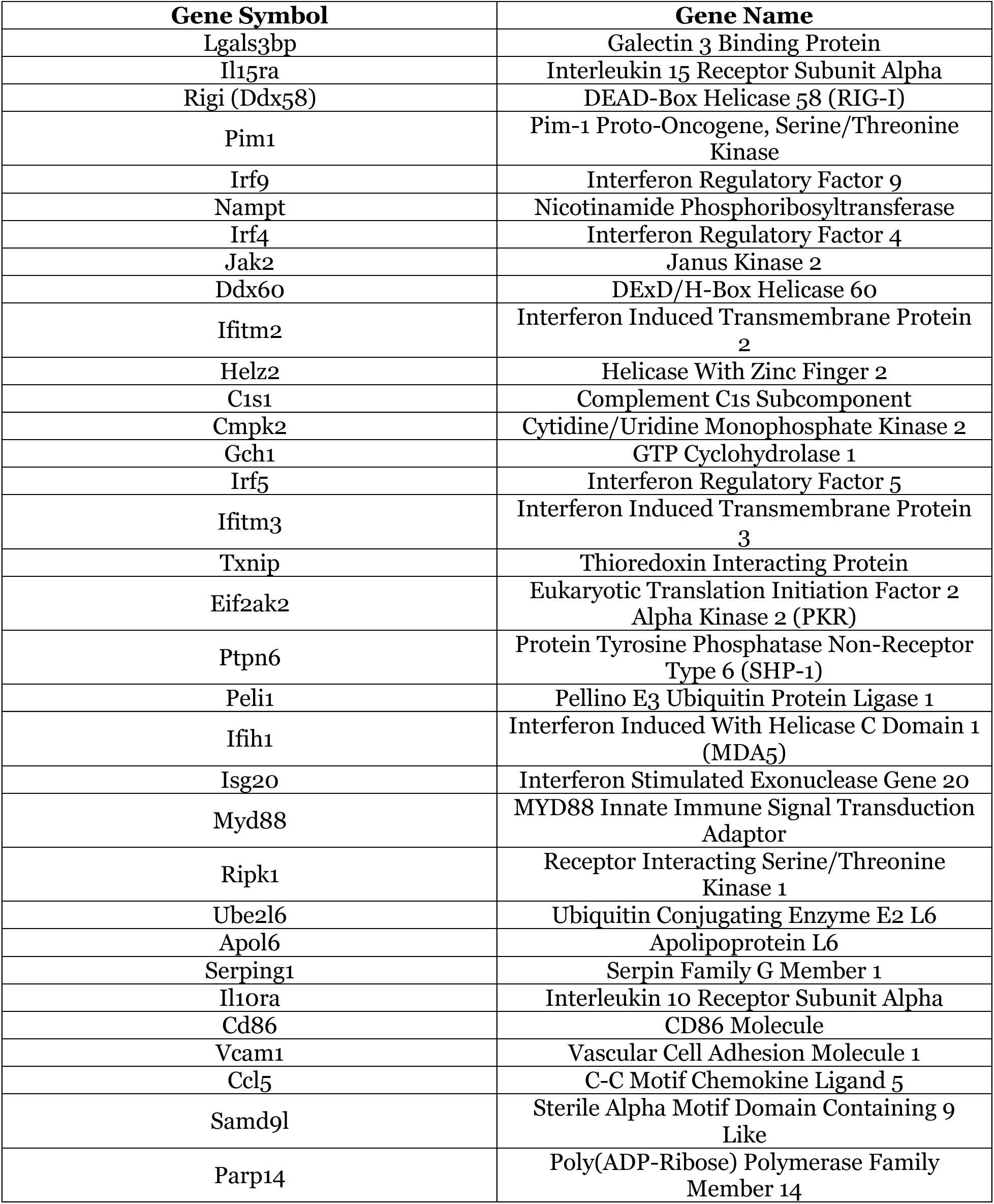
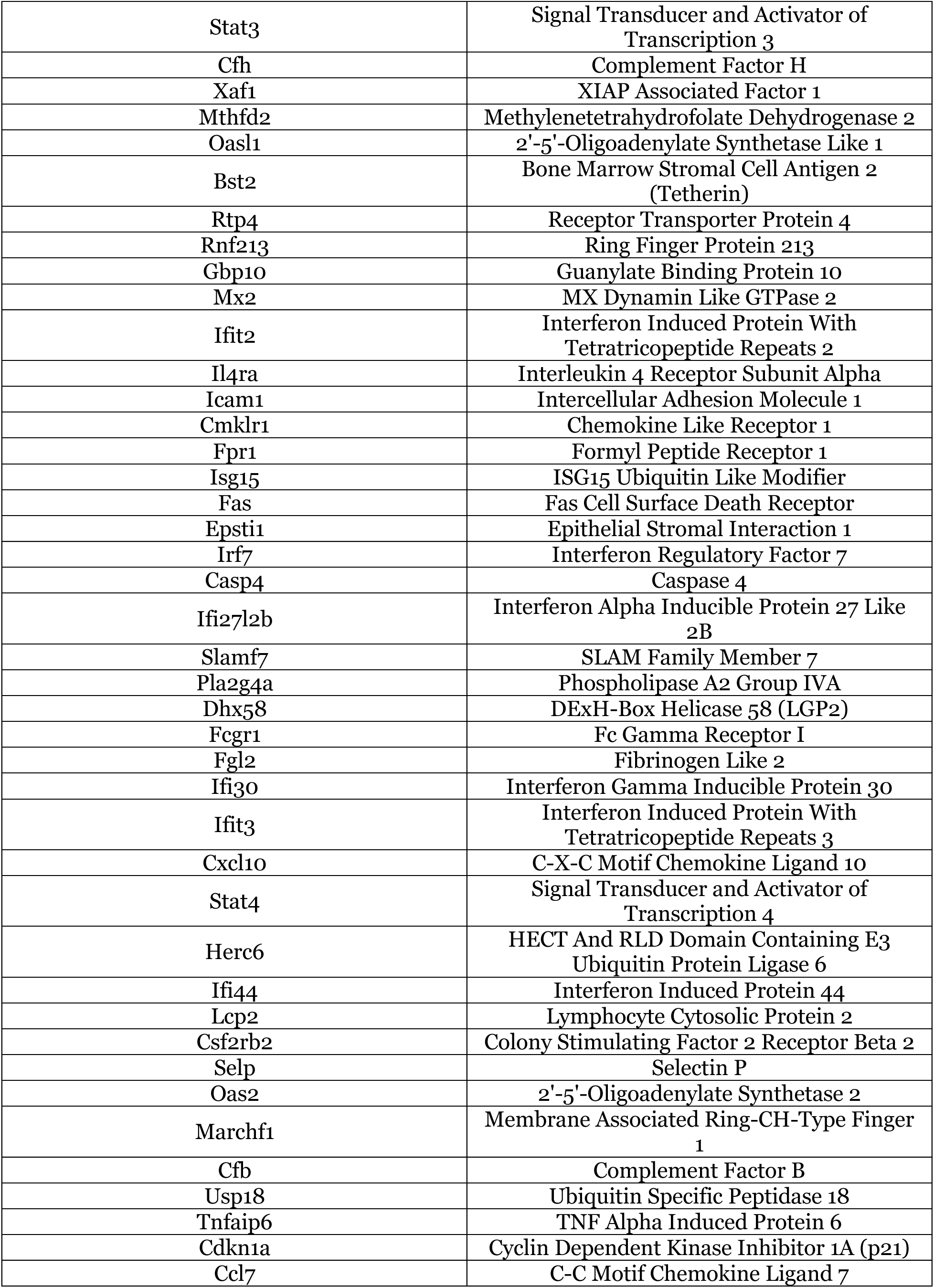
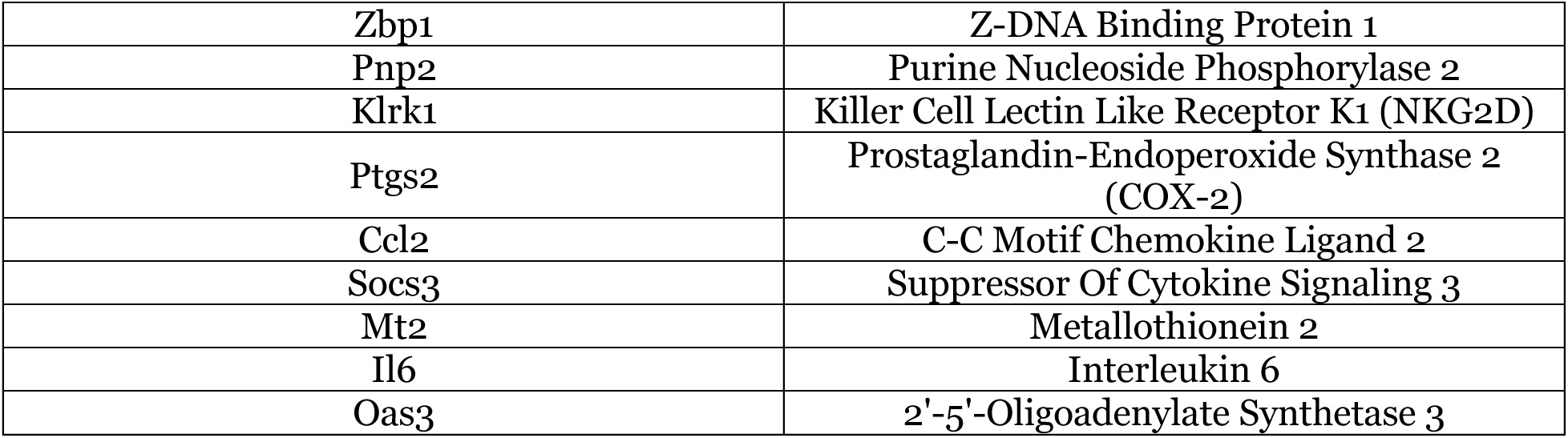
Suppression of the Hallmark IFNγ Response gene set in aged p38β^-/-^ hearts. The Hallmark IFNγ Response gene set was identified by GSEA as one of the top 10 suppressed gene sets in the hearts of aged p38β^-/-^ male mice compared with WT counterparts. The data in this table are related to Figure 8J.

**Supplemental Table 8.**
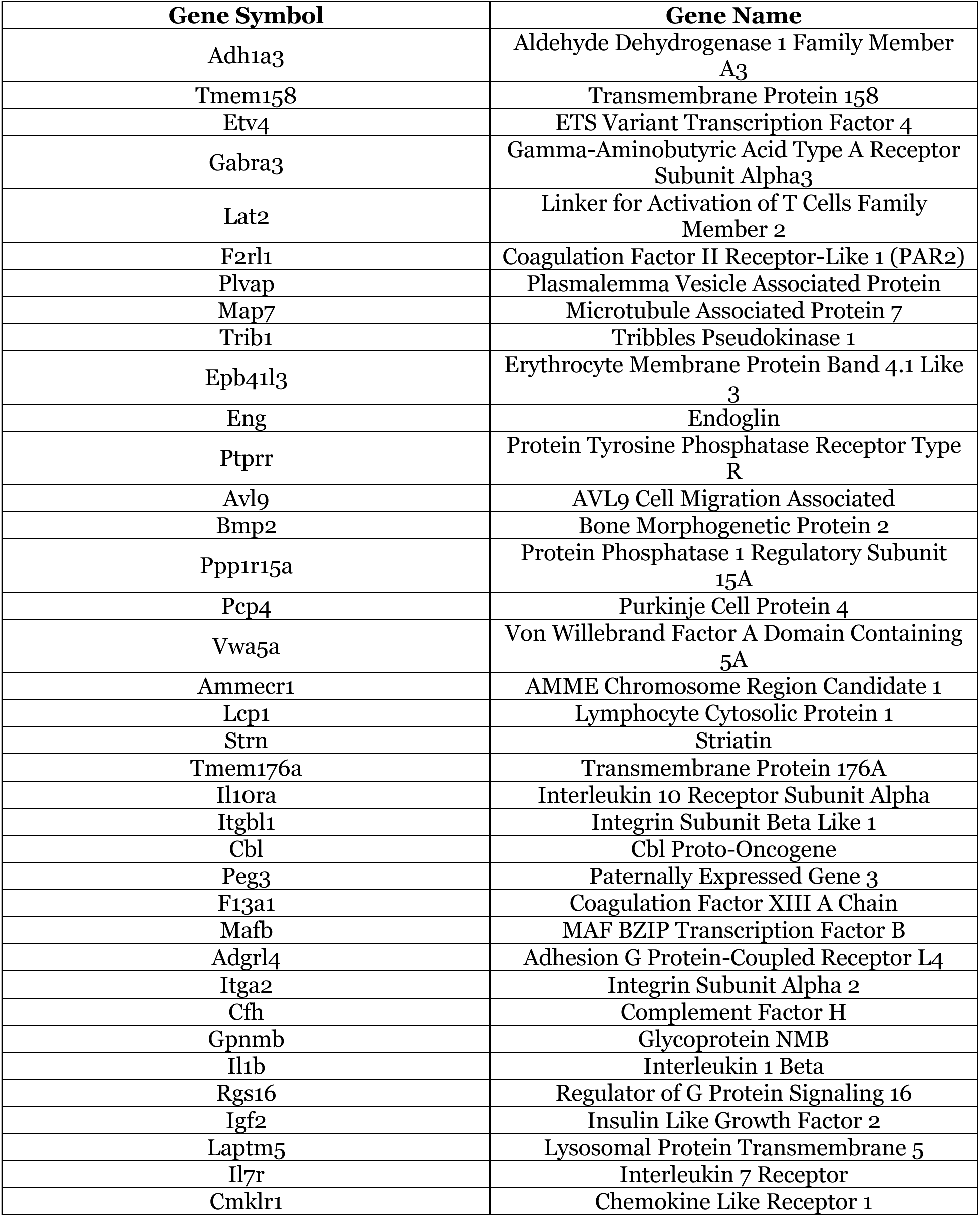
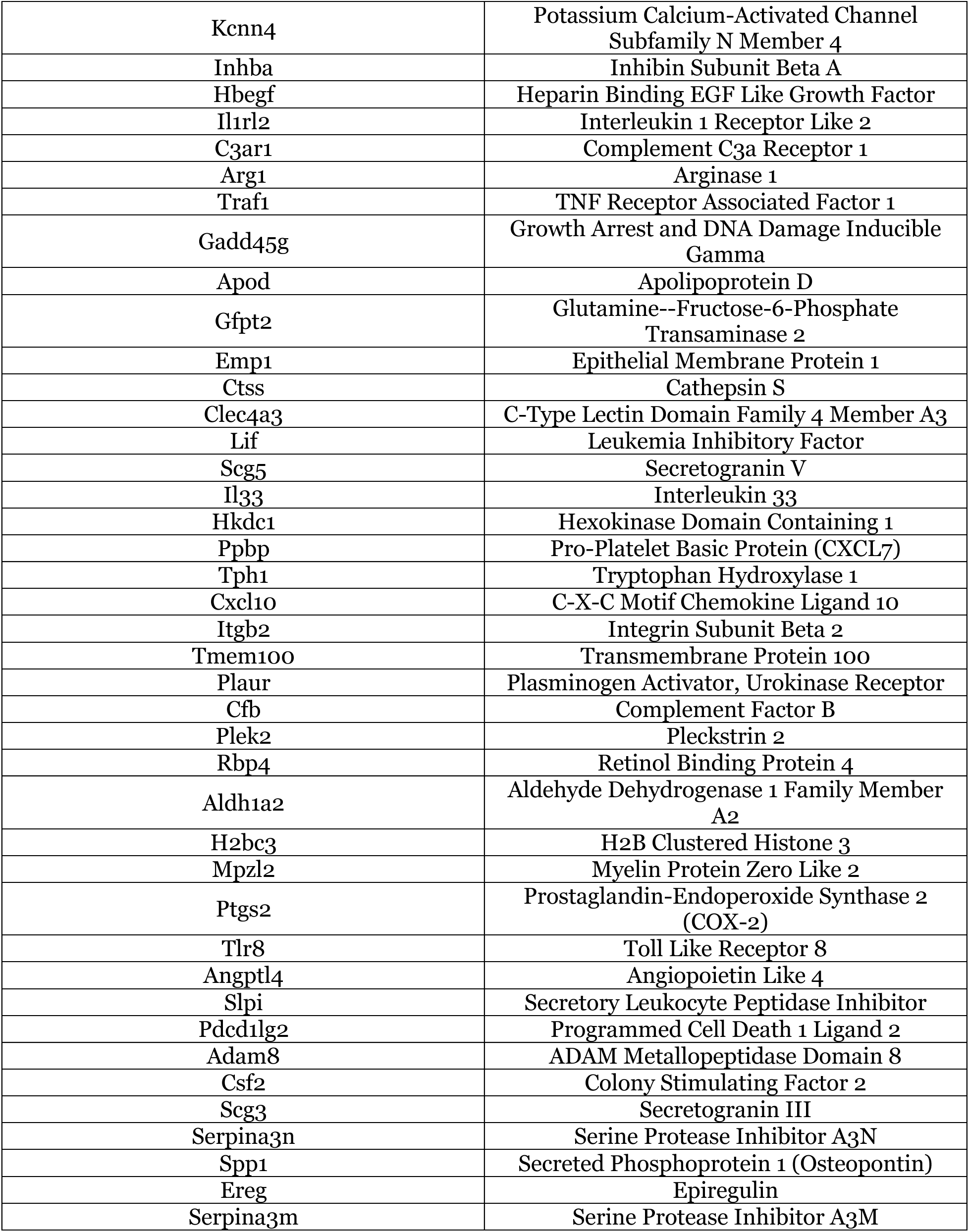
Suppression of the Hallmark Kras Signaling Up gene set in aged p38β^-/-^ hearts. The Hallmark Kras Signaling Up gene set was identified by GSEA as one of the top 10 suppressed gene sets in the hearts of aged p38β^-/-^ male mice compared with WT counterparts. The data in this table are related to Figure 8J.

**Supplemental Table 9.**
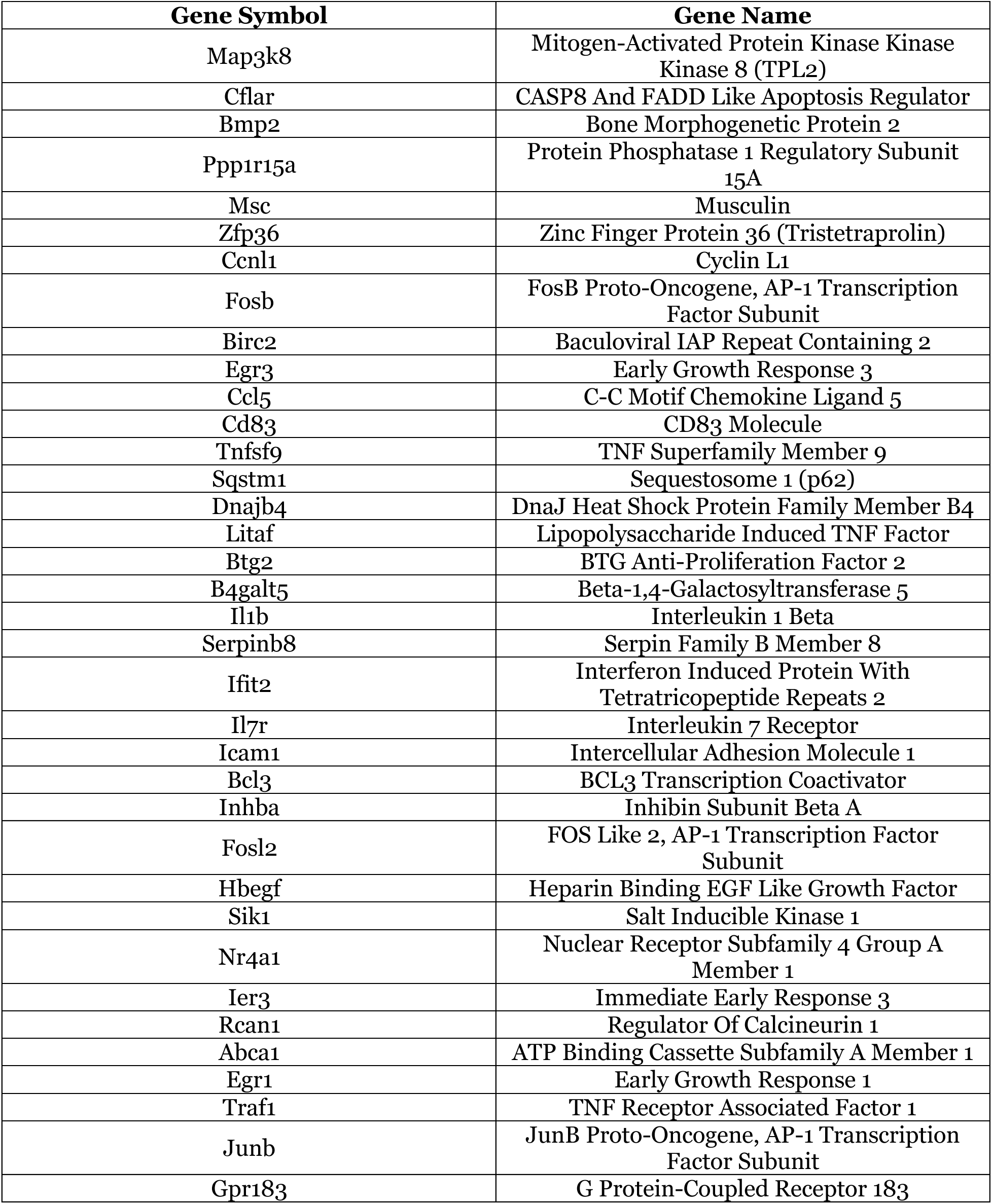
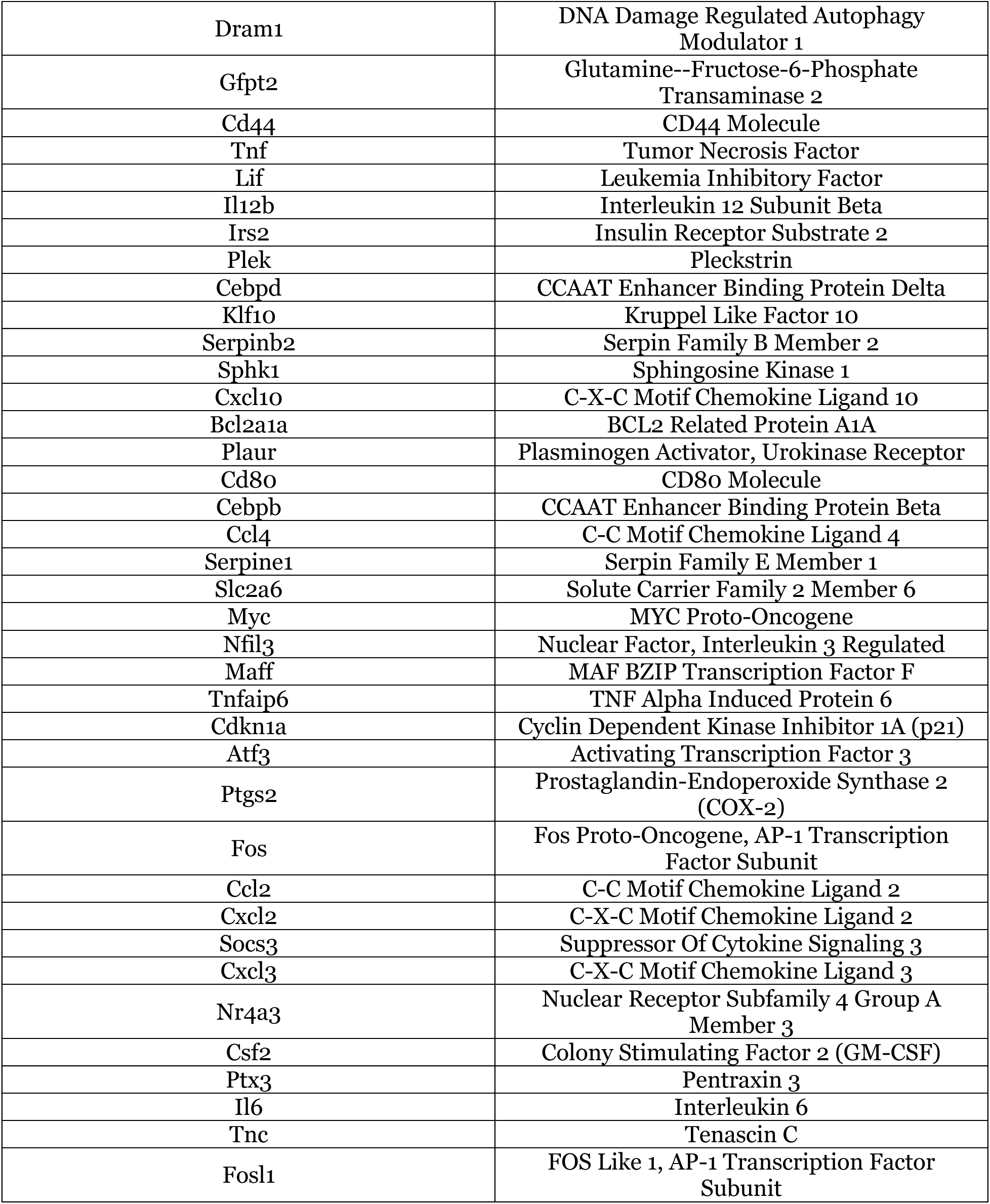
Suppression of the Hallmark TNFα Signaling via NFKB gene set in aged p38β^-/-^ hearts. The Hallmark TNFα Signaling via NFKB gene set was identified by GSEA as one of the top 10 suppressed gene sets in the hearts of aged p38β^-/-^ male mice compared with WT counterparts. The data in this table are related to Figure 8J.

**Supplemental Table 10.** Suppression of the Hallmark Inflammatory Response gene set in aged p38β^-/-^ hearts. The Hallmark Inflammatory Response gene set was identified by GSEA as one of the top 10 suppressed gene sets in the hearts of aged p38β^-/-^ male mice compared with WT counterparts. The data in this table are related to Figure 8J.

| Gene Symbol | Gene Name |
| --- | --- |
| Ccl5 | C-C Motif Chemokine Ligand 5 |
| Tnfsf9 | TNF Superfamily Member 9 |
| Rnf144b | Ring Finger Protein 144B |
| Btg2 | BTG Anti-Proliferation Factor 2 |
| Il1b | Interleukin 1 Beta |
| Mefv | Mediterranean Fever Gene (Pyrin) |
| Bst2 | Bone Marrow Stromal Cell Antigen 2 |
| Rgs16 | Regulator of G Protein Signaling 16 |
| Rtp4 | Receptor Transporter Protein 4 |
| Il7r | Interleukin 7 Receptor |
| Il4ra | Interleukin 4 Receptor Subunit Alpha |
| Icam1 | Intercellular Adhesion Molecule 1 |
| Csf3r | Colony Stimulating Factor 3 Receptor |
| Cmklr1 | Chemokine Like Receptor 1 |
| Fpr1 | Formyl Peptide Receptor 1 |
| Has2 | Hyaluronan Synthase 2 |
| Inhba | Inhibin Subunit Beta A |
| Hbegf | Heparin Binding EGF Like Growth Factor |
| Adora2b | Adenosine A2b Receptor |
| Abca1 | ATP Binding Cassette Subfamily A Member 1 |
| C5ar1 | Complement C5a Receptor 1 |
| Itgb8 | Integrin Subunit Beta 8 |
| Cd55 | CD55 Molecule (Decay Accelerating Factor) |
| Pdpn | Podoplanin |
| C3ar1 | Complement C3a Receptor 1 |
| Irf7 | Interferon Regulatory Factor 7 |
| Ifitm1 | Interferon Induced Transmembrane Protein 1 |
| Msr1 | Macrophage Scavenger Receptor 1 |
| Gpr183 | G Protein-Coupled Receptor 183 |
| Pik3r5 | Phosphoinositide-3-Kinase Regulatory Subunit 5 |
| Itgb3 | Integrin Subunit Beta 3 |
| Lif | Leukemia Inhibitory Factor |
| Osmr | Oncostatin M Receptor |
| Met | MET Proto-Oncogene, Receptor Tyrosine Kinase |
| Il12b | Interleukin 12 Subunit Beta |
| Aqp9 | Aquaporin 9 |
| Csf3 | Colony Stimulating Factor 3 |
| Nmur1 | Neuromedin U Receptor 1 |
| Cd14 | CD14 Molecule |
| Marco | Macrophage Receptor With Collagenous Structure |
| Sphk1 | Sphingosine Kinase 1 |
| Cxcl10 | C-X-C Motif Chemokine Ligand 10 |
| Lcp2 | Lymphocyte Cytosolic Protein 2 |
| Vip | Vasoactive Intestinal Peptide |
| Sgms2 | Sphingomyelin Synthase 2 |
| Plaur | Plasminogen Activator, Urokinase Receptor |
| Serpine1 | Serpin Family E Member 1 (PAI-1) |
| Myc | MYC Proto-Oncogene |
| Ccl22 | C-C Motif Chemokine Ligand 22 |
| Tpbg | Trophoblast Glycoprotein |
| Ccr7 | C-C Motif Chemokine Receptor 7 |
| Tnfaip6 | TNF Alpha Induced Protein 6 |
| Slc1a2 | Solute Carrier Family 1 Member 2 |
| Ccl17 | C-C Motif Chemokine Ligand 17 |
| Il18rap | Interleukin 18 Receptor Accessory Protein |
| Kcna3 | Potassium Voltage-Gated Channel Subfamily A Member 3 |
| Cdkn1a | Cyclin Dependent Kinase Inhibitor 1A (p21) |
| Ccl7 | C-C Motif Chemokine Ligand 7 |
| Osm | Oncostatin M |
| Tnfsf15 | TNF Superfamily Member 15 |
| Il10 | Interleukin 10 |
| Il18r1 | Interleukin 18 Receptor 1 |
| Ccl2 | C-C Motif Chemokine Ligand 2 |
| Rgs1 | Regulator of G Protein Signaling 1 |
| Timp1 | TIMP Metalloproteinase Inhibitor 1 |
| Il6 | Interleukin 6 |
| Bdkrb1 | Bradykinin Receptor B1 |
| Ereg | Epiregulin |

**Supplemental Table 11.**
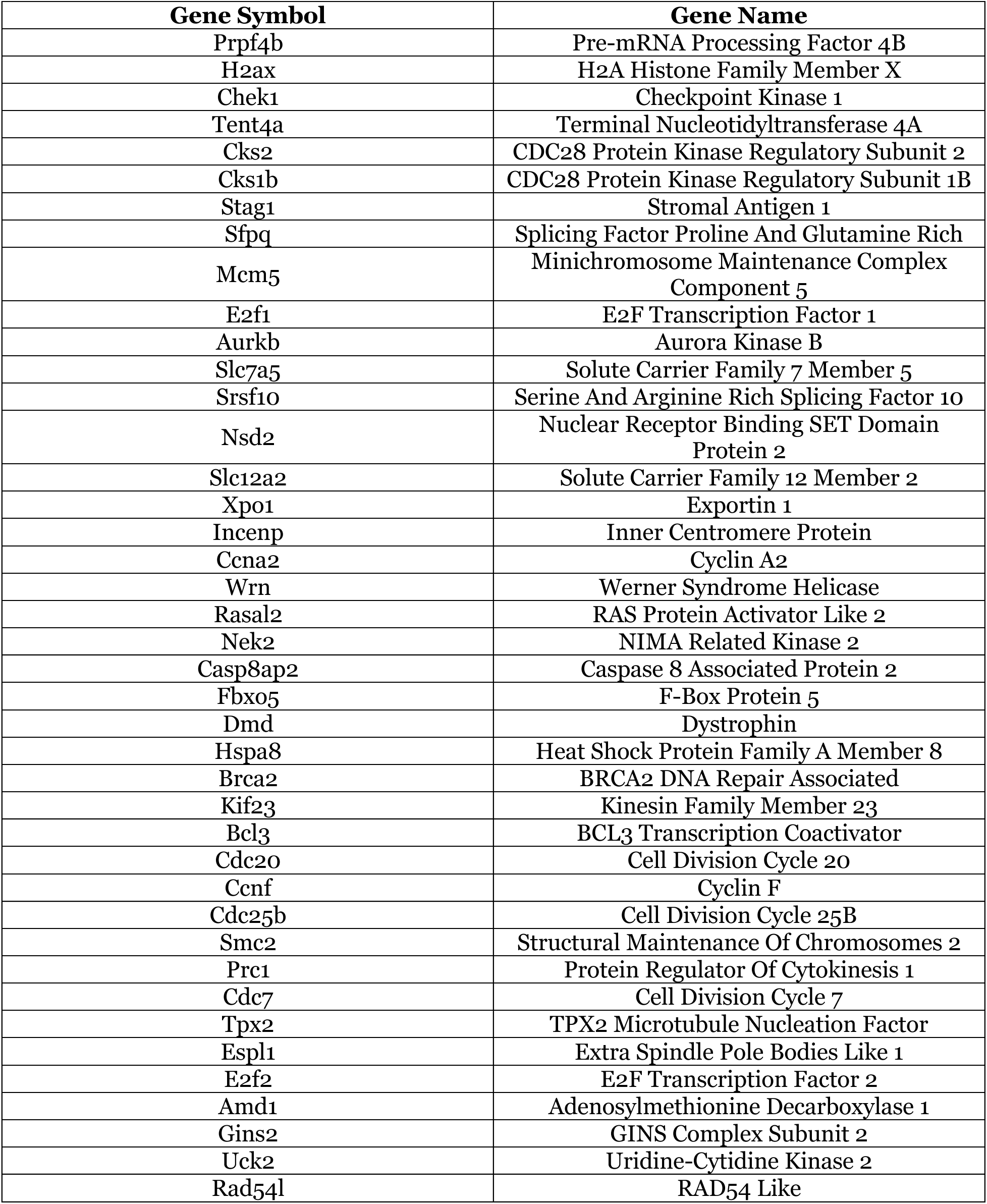
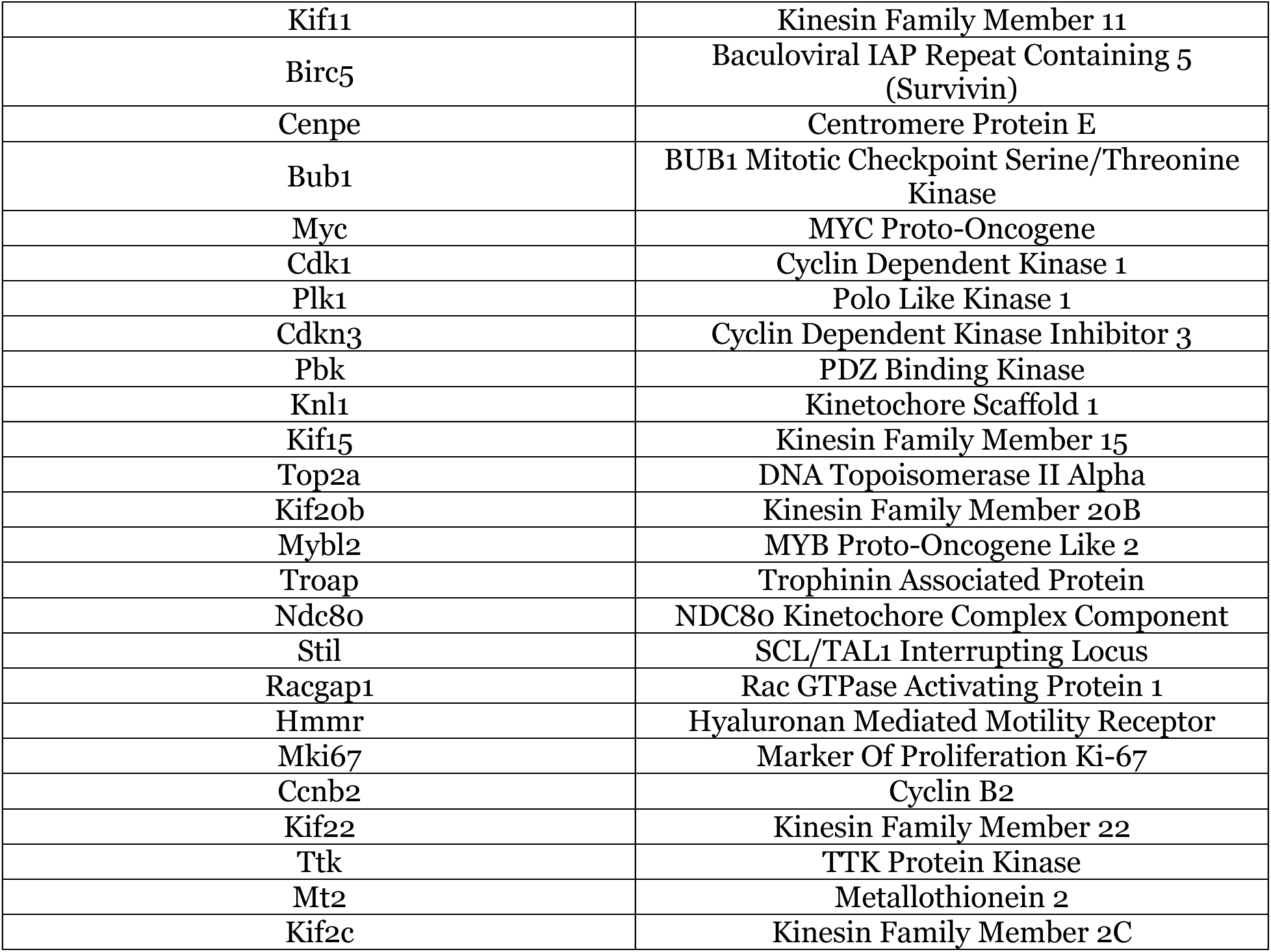
Suppression of the Hallmark G2M Checkpoint gene set in aged p38β^-/-^ hearts. The Hallmark G2M Checkpoint gene set was identified by GSEA as one of the top 10 suppressed gene sets in the hearts of aged p38β^-/-^ male mice compared with WT counterparts. The data in this table are related to Figure 8J.

**Supplemental Table 12.**
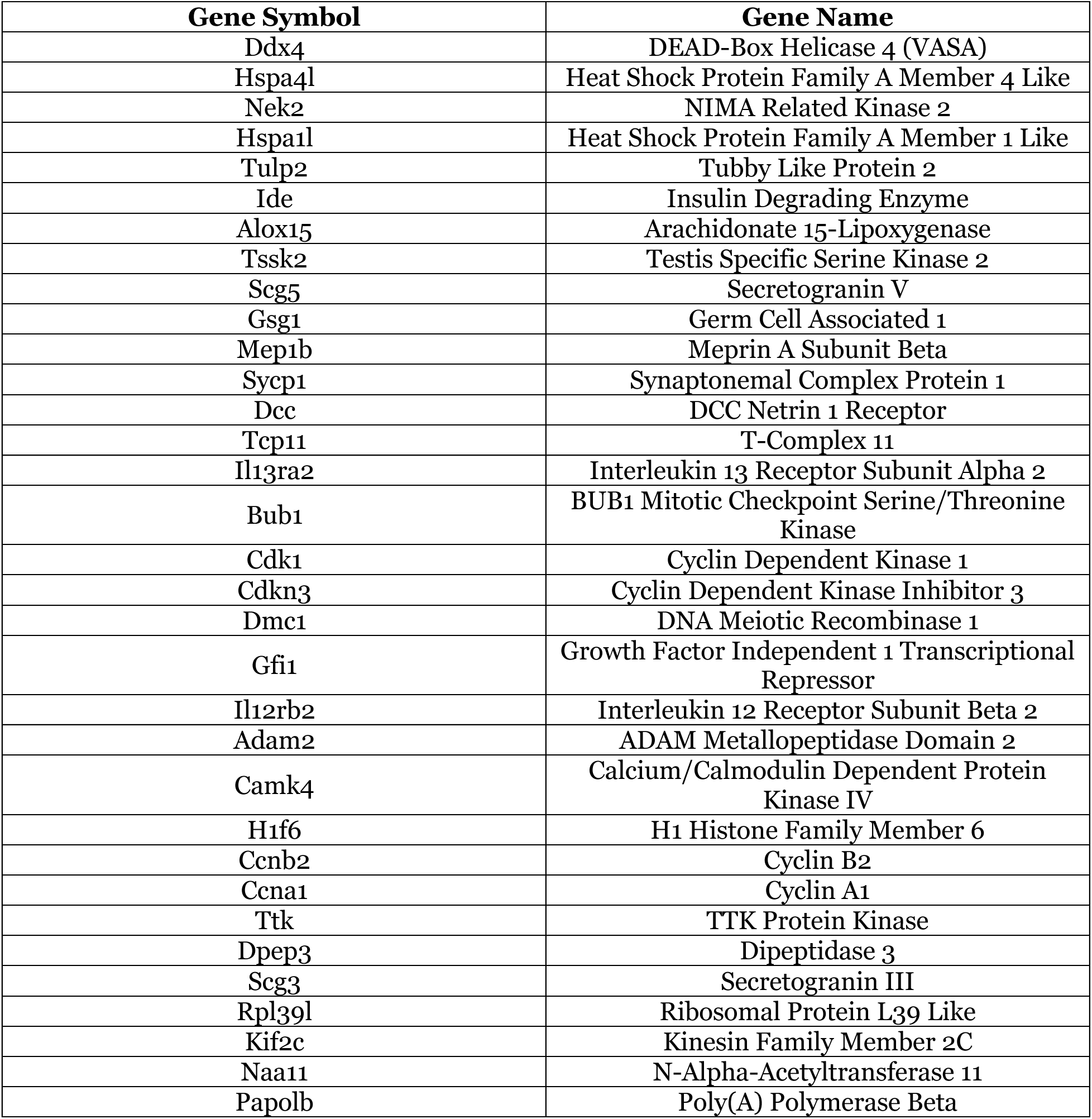
Suppression of the Hallmark Spermatogenesis gene set in aged p38β^-/-^ hearts. The Hallmark Spermatogenesis gene set was identified by GSEA as one of the top 10 suppressed gene sets in the hearts of aged p38β^-/-^ male mice compared with WT counterparts. The data in this table are related to Figure 8J.

**Supplemental Table 13.**
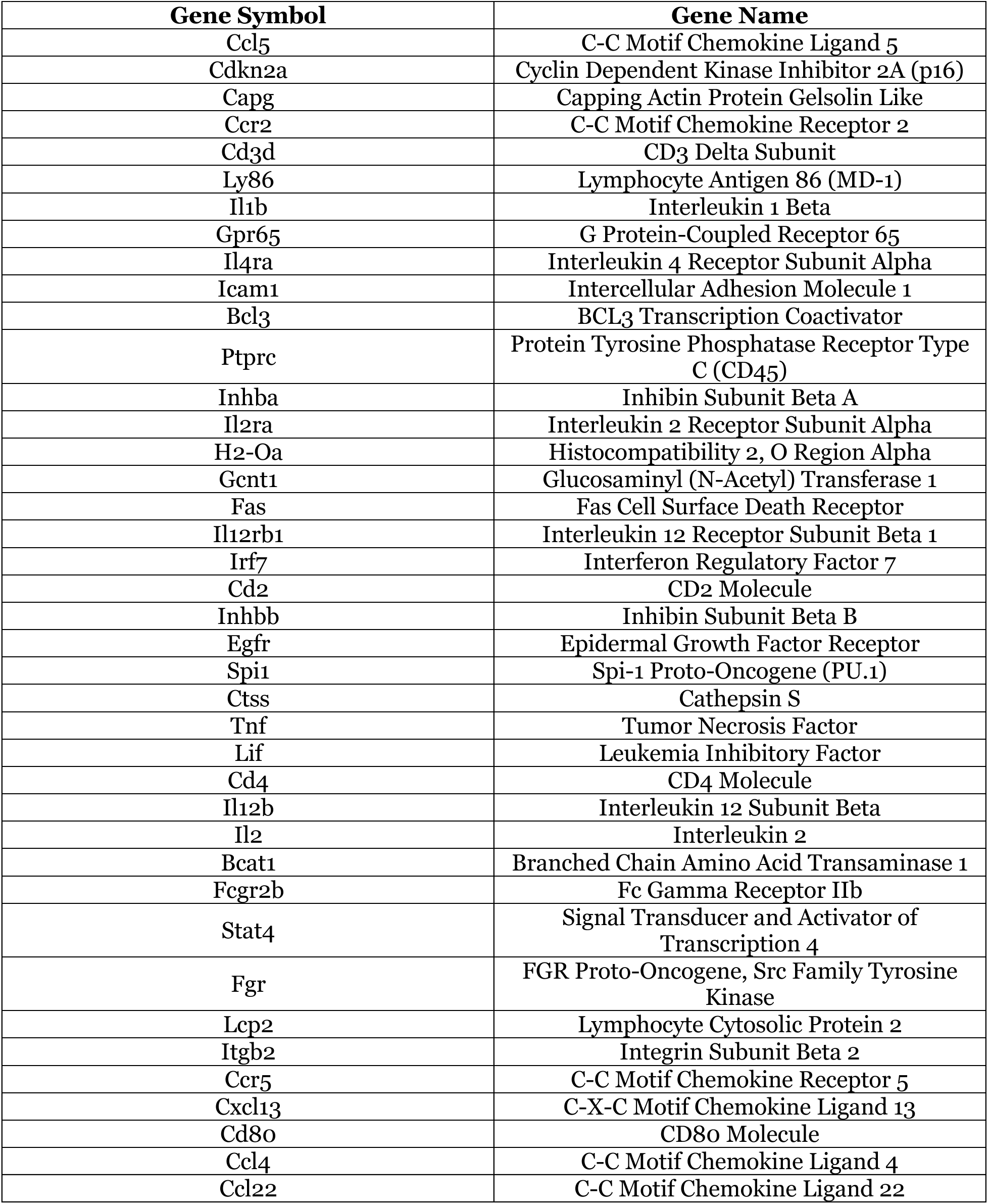
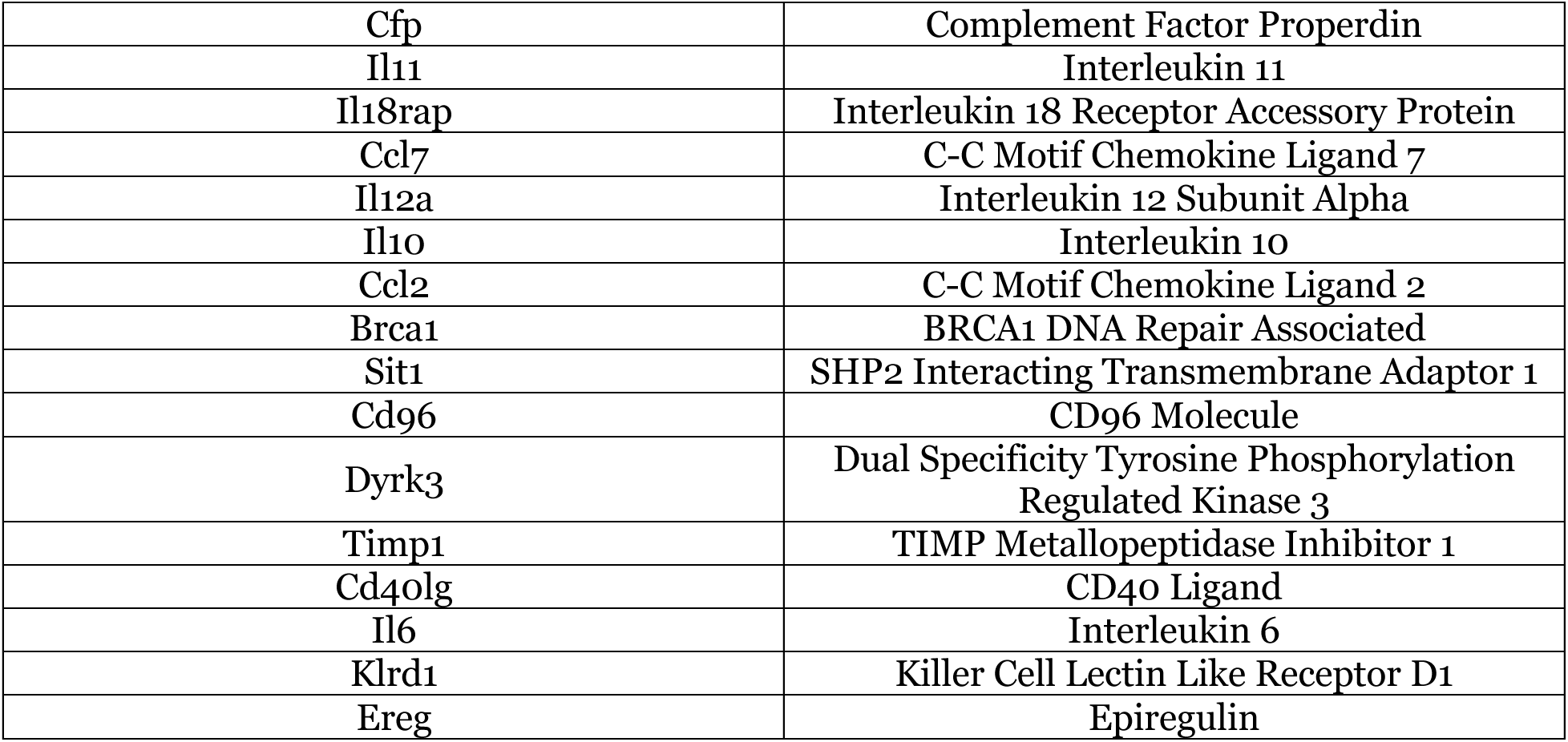
Suppression of the Hallmark Allograft Rejection gene set in aged p38β^-/-^ hearts. The Hallmark Allograft Rejection gene set was identified by GSEA as one of the top 10 suppressed gene sets in the hearts of aged p38β^-/-^ male mice compared with WT counterparts. The data in this table are related to Figure 8J.

**Supplemental Table 14.**
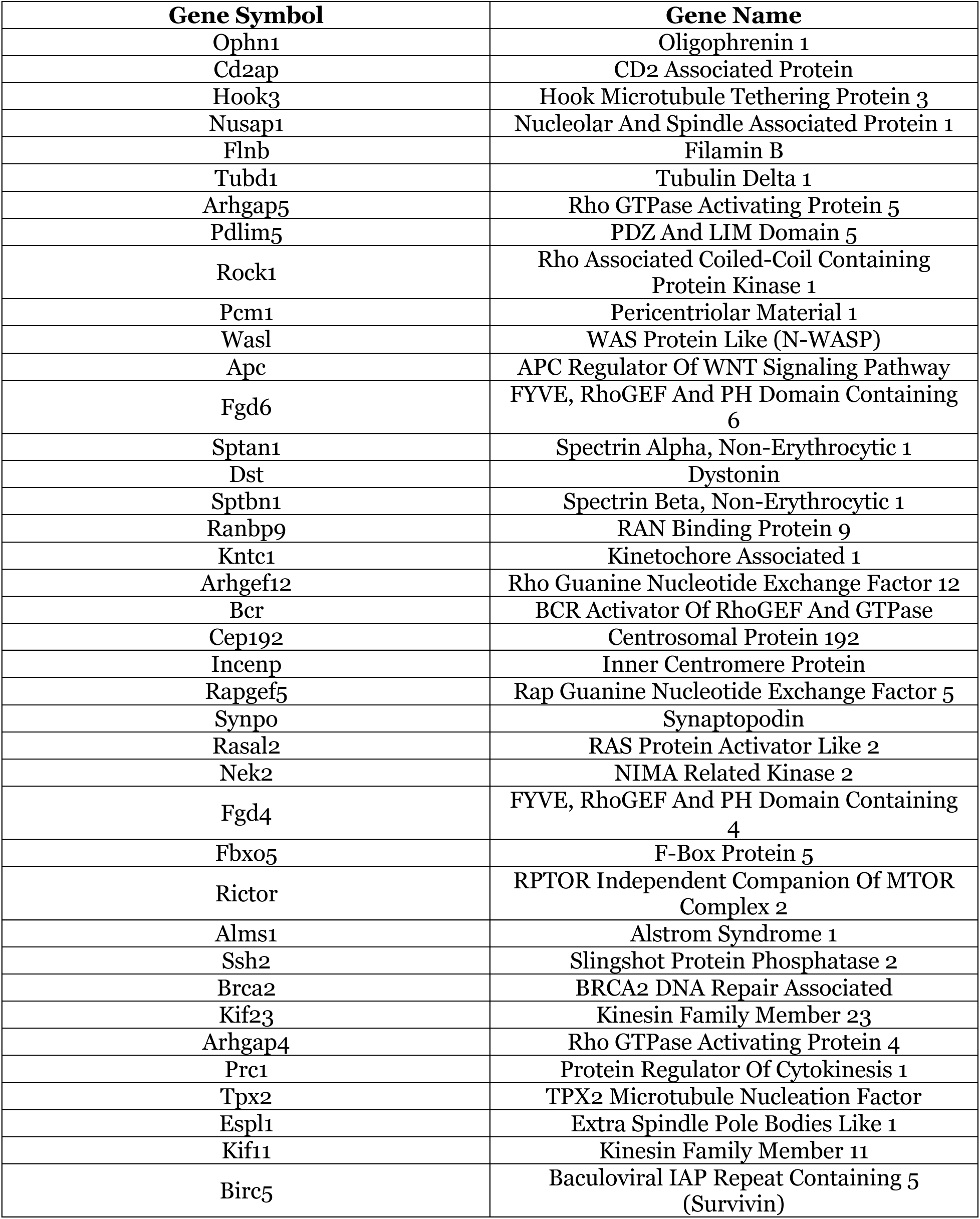
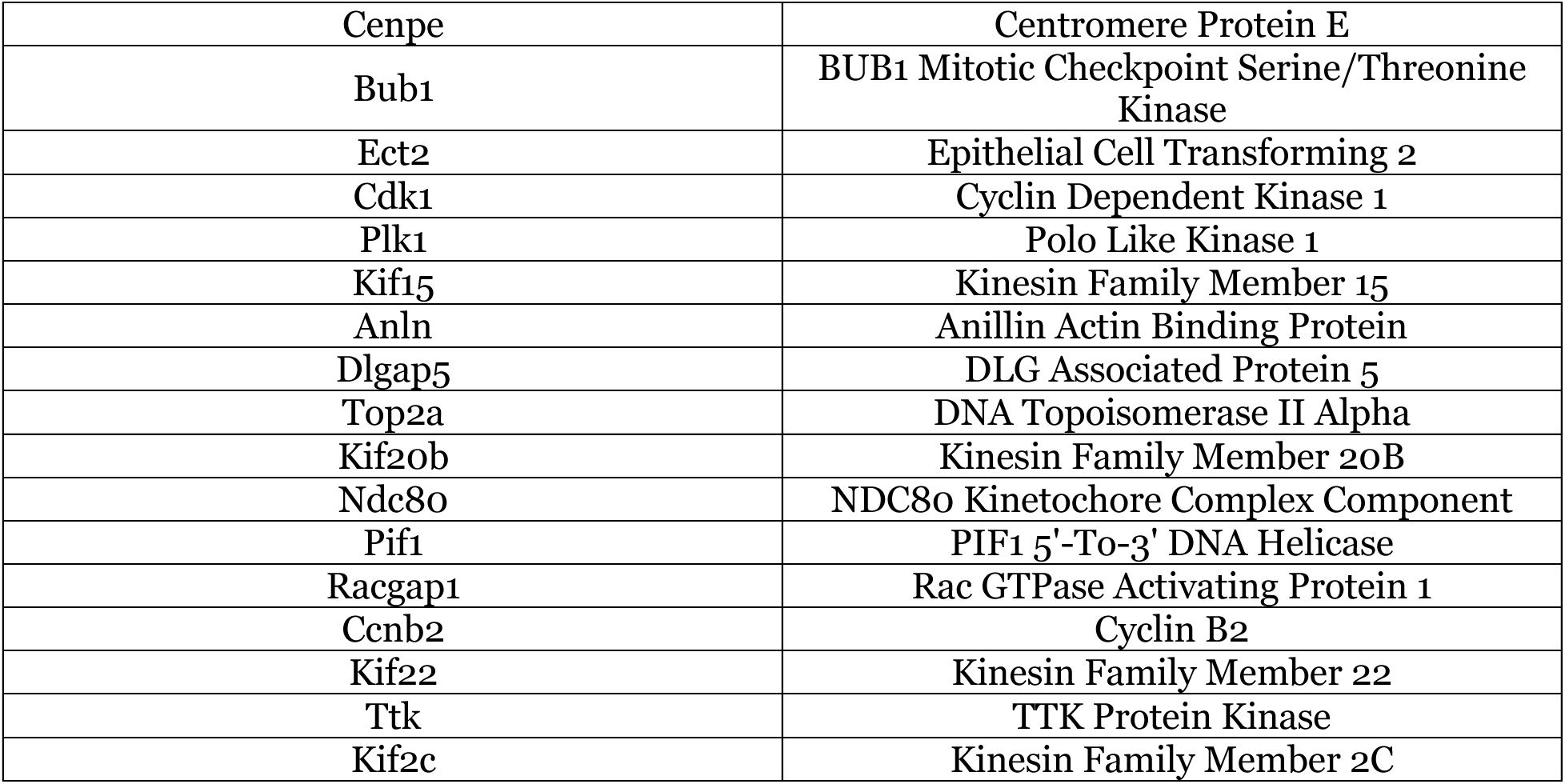
Suppression of the Hallmark Mitotic Spindle gene set in aged p38β^-/-^ hearts. The Hallmark Mitotic Spindle gene set was identified by GSEA as one of the top 10 suppressed gene sets in the hearts of aged p38β^-/-^ male mice compared with WT counterparts. The data in this table are related to Figure 8J.

**Supplemental Figure 1.**
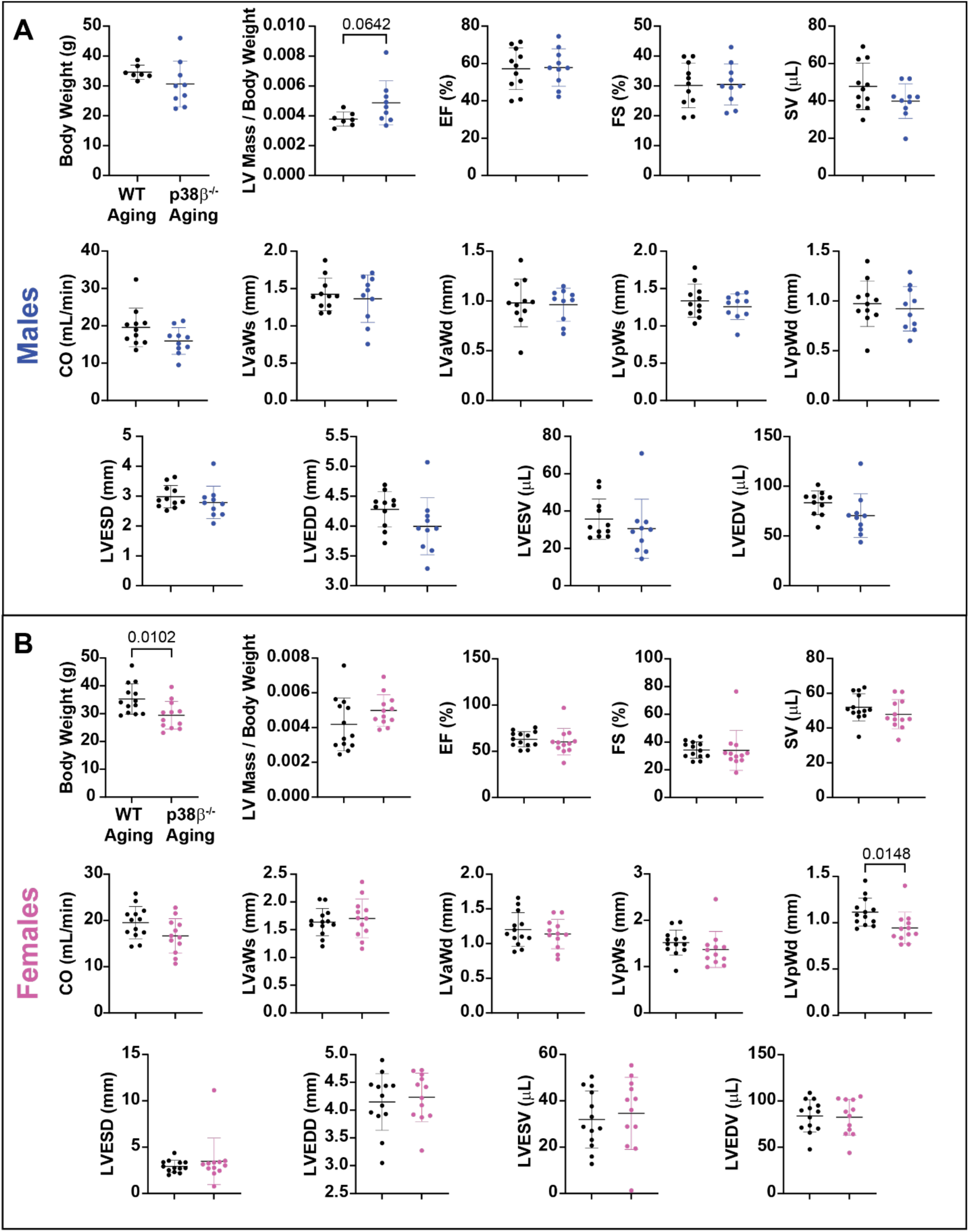
Sex-stratified analysis of cardiac structural and functional remodeling in aged WT and p38β^-/-^ mice. LV short-axis echocardiogram parameter quantification in aged WT and p38β^-/-^ male (***A***) and female (***B***) mice. LV anterior and posterior wall thicknesses during systole and diastole (LVaWs, LVaWd, LVpWs, LVpWd), LV end-systolic diameter (LVESD), LV end-diastolic diameter (LVEDD), and LV volume during systole and diastole (LVESV and LVEDV). Sample sizes: WT males: n = 11; p38β^-/-^ males: n = 9; WT females: n = 10; p38β^-/-^ females: n = 9.

**Supplemental Figure 2.**
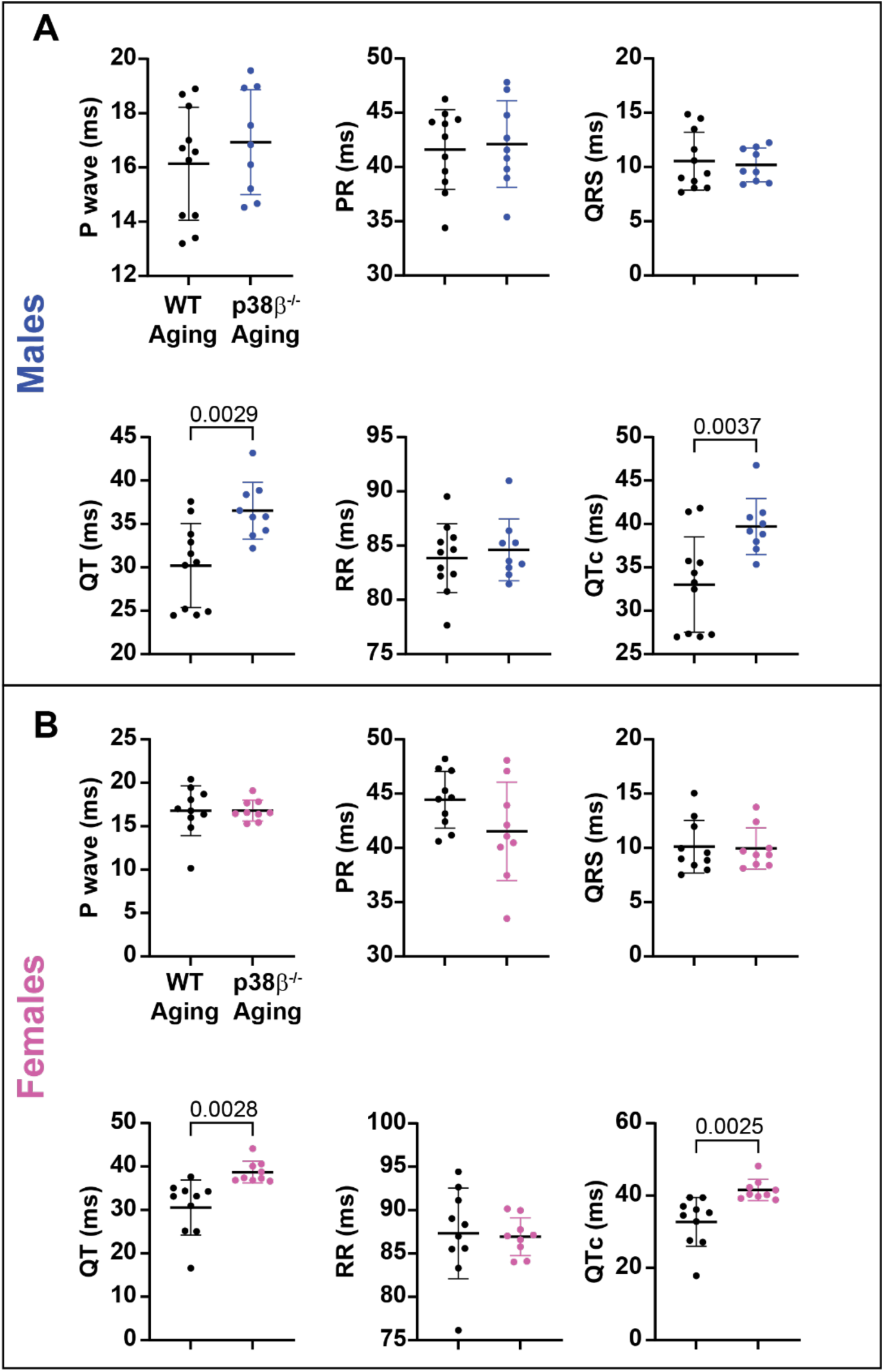
p38β deficiency prolongs QT and QTc in aged male and female mice. ECG parameters were analyzed in aged male (***A***) and female (**B**) mice. Unpaired Student’s *t*-tests were used to assess statistical significance between groups. Sample sizes: WT males: n = 11; p38β^-/-^ males: n = 9; WT females: n = 10; p38β^-/-^ females: n = 9.

**Supplemental Figure 3.**
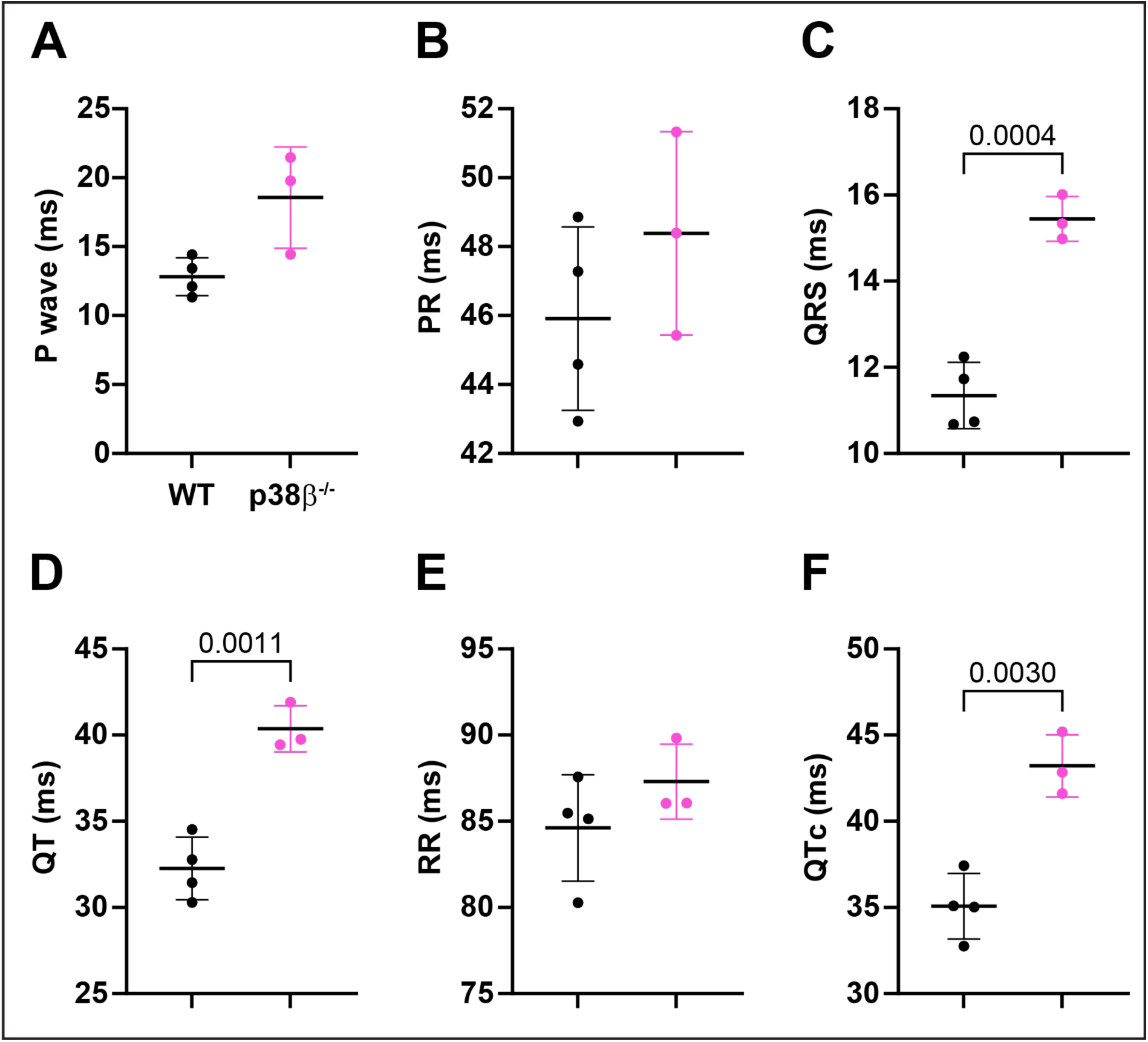
p38β deficiency prolongs QT and QTc in 20-month-old female mice. Quantification of ECG waveform parameters, including P wave duration (***A***), PR interval (***B***), QRS interval (***C***), QT interval (***D***), RR interval (***E***), and rate-corrected QTc interval (***F***) in 20-month-old WT and p38β^-/-^ female mice. Unpaired Student’s *t*-tests were used to assess statistical significance between groups. Sample sizes: WT females: n = 4; p38β^-/-^ females: n = 3.

